# The DNA-repair protein APE1 participates with hnRNPA2B1 to motif-enriched and prognostic miRNA secretion

**DOI:** 10.1101/2024.02.02.578563

**Authors:** Giovanna Mangiapane, Michela Notarangelo, Giulia Canarutto, Fabrizio Fabbiano, Emiliano Dalla, Monica Degrassi, Giulia Antoniali, Nicolò Gualandi, Veronica De Sanctis, Silvano Piazza, Vito Giuseppe D’Agostino, Gianluca Tell

## Abstract

The base excision repair (BER) Apurinic/apyrimidinic endonuclease 1 (APE1) enzyme is endowed with several non-repair activities including miRNAs processing. APE1 is overexpressed in many cancers but its causal role in the tumorigenic processes is largely unknown. We recently described that APE1 can be actively secreted by mammalian cells through exosomes. However, APE1 role in EVs or exosomes is still unknown, especially regarding a putative regulatory function on small non-coding RNAs vesicular secretion. Through dedicated transcriptomic analysis on cellular and vesicular small RNAs of different APE1-depleted cancer cell lines, we found that miRNAs loading into EVs is a regulated process, dependent on APE1, distinctly conveying RNA subsets into vesicles. We identified APE1-dependent secreted miRNAs characterized by enriched sequence motifs and possible binding sites for APE1. In 33 out of 34 APE1-dependent-miRNA precursors, we surprisingly found EXO-motifs and proved that APE1 cooperates with hnRNPA2B1 for the EV-sorting of a subset of miRNAs, including miR-1246, through direct binding to GGAG stretches. Using TCGA-datasets, we showed that these miRNAs identify a signature with high prognostic significance in cancer. In summary, we provided evidence that APE1 is part of the protein cargo of secreted EVs, suggesting a novel post-transcriptional role for this ubiquitous DNA-repair enzyme that could explain its role in cancer progression.

## INTRODUCTION

Apurinic/apyrimidinic endodeoxyribonuclease 1 (APE1) is the main AP-endodeoxyribonuclease belonging to the base excision repair (BER) pathway. Once alkylating agents or oxidative stress create DNA lesions, APE1 recognizes and cleaves at the 5’ of the deoxy-ribose (dRP)-abasic site generated by glycosylases, such as OGG1 (1). Besides this canonical role, APE1 recently emerged as a relevant factor in the adaptive cellular response to genotoxic stress, regulating a plethora of coding and non-coding RNAs (ncRNAs) (2, 3) and the interplay required for gene expression (4). Interestingly, APE1 also shows endoribonuclease activity on abasic sites present in single-stranded RNAs (5), whose origin is still debated (6). We found that APE1 is involved in the processing of rAP and r8oxoG both in RNA and DNA (7, 8), thus representing a novel component of the damaged-RNA cleansing machinery (2, 9).

The structure of APE1 is characterized by three different functional sub-domains, which are responsible for the multifunctional roles of the protein: i) The unstructured N-terminal region (from aa 1 to aa 35/37), which stabilizes protein-protein- or protein-RNA-interactions (10); ii) The central region (from aa 35 to aa 127), responsible for the “redox mechanism” involved in gene transcription, and iii) The C-terminal region, which exerts the endodeoxyribonuclease activity (11).

An extended analysis of the APE1-interactome network revealed proteins belonging to five biological pathways with the following representativity: RNA processing, DNA Repair, double-strand breaks (DSB) repair, transcriptional regulation, and apoptosis (12, 13), suggesting that the interaction with different partners could possibly direct a spatial-temporal activity of APE1, working as a molecular switch to exert its functions on different substrates (13). The pivotal role of APE1 in RNA metabolism is indicated by 60% of interacting partners devoted to RNA-processing and RNA-binding activities (12), which we and others started to elucidate for microRNAs (miRNAs) (12, 13). miRNAs are a class of small ncRNAs that post-transcriptionally regulate gene expression (14). Mature miRNAs are short sequences of about 22 nucleotides, which pair with target mRNAs’ 3′-untranslated regions (UTRs) to inhibit their translation and/or promote their degradation (15). The activity of miRNAs is relevant in physiological and pathological conditions, including cell metabolism, cardiovascular diseases, and hallmarks of cancer spanning from cell proliferation to tumor progression and chemoresistance (14, 16). We demonstrated that, upon genotoxic stress, nuclear APE1 favors the processing and stability of some miRNA precursors through the association with the DROSHA microprocessor complex, impacting, for example, on the miR-221/222 axis which, in turn, modulates the expression of the tumor suppressor PTEN (12). In addition, we also showed that reduced levels of APE1 compromise the expression of oncogenic miRNAs, impacting the epidermal-mesenchymal transition process, in Non-Small cell Lung Cancer (NSCLC) models, through DICER expression (17).

A fascinating new field of research relies on the unexpected finding that APE1 can be secreted in the extracellular milieu. Notably, relatively high APE1 blood levels were associated with tumor progression and chemoresistance, suggesting that serum APE1 (sAPE1) could work as a predictive biomarker in solid tumors, such as NSCLC, bladder cancer, and hepatocellular carcinoma (HCC) (18–21). We recently found that APE1 can be detected in extracellular vesicles (EVs), including exosomes, released by mammalian cells (22). EVs are nano/submicro-sized lipid particles involved in cell-to-cell communication (23). They shuttle a variety of RNA species, not limited to ncRNAs, miRNAs, and mRNAs, that can be functionally transferred to the recipient cells (24) through still un-deciphered paracrine mechanisms (25). In the context of the tumor microenvironment, EVs represent vehicles of factors promoting cancer progression (26). Therefore, the study of the mechanisms involved in sorting specific EV-RNA molecules can be extraordinarily advantageous to disentangle the complexity of the mechanisms activated by cancer cells (27), offering opportunities for developing novel diagnostic and/or drug-targeting strategies.

Among the RNA molecules present in EVs or exosomes, miRNAs represent hubs of processing, sorting, and quality control steps, likely resulting from a selective process (28) since not all the intracellular miRNA heterogeneity seems indistinctly conveyed into EVs (29) or independent from trans-acting factors (30, 31). For instance, nucleophosmin (NPM1) prevents the degradation of microRNAs within exosomes (32). On the other hand, RNA binding proteins (RBPs) such as ELAV like RNA-binding protein 1 (ELAVL1), Y-box protein 1 (YBX1), or especially several heterogeneous nuclear ribonucleoproteins (hnRNP) family members, may recognize specific consensus sequences (known as EXO-motifs) (24, 32) associated with a preferential EV-packaging (31). By virtue of the connections between APE1 activity and consequent miRNA regulation, knowing that exosomal APE1 is enzymatically active and promptly secreted in the presence of genotoxic stress (22), in this work we investigated the direct involvement of APE1 in EV-miRNA sorting pathways. Through integrative cellular, biochemical, and computational approaches, we here proved evidence that APE1 cooperates with hnRNPA2B1 in selecting vesicular miRNAs characterized by hallmark cis-elements, providing the first evidence of a DNA repair enzyme participating in the secretion of miRNA subsets potentially relevant for disease interception.

## RESULTS

### APE1 regulates the steady-state levels of intracellular- and EV-sorted RNA abundance

Given the emerging role of APE1 in RNA processing and its presence into EVs (22), we speculated that APE1 might participate in vesicular RNA sorting mechanisms. To test this hypothesis, we employed a quantitative approach to investigate the relative abundance of RNA present in vesicles released from CH12F3 lymphoma cell lines already characterized for expressing or not APE1 (CH12F3Δ/+/+ and CH12F3Δ/Δ/Δ cells, respectively) (33). We used nickel-based isolation (NBI) procedure for collecting polydisperse EVs and applying RNA analysis, according to previous literature (34, 35). We subjected the serum-free conditioned media, after 24 hr exposure to the cells, to NBI. Then, we characterized the isolated particles by tunable resistive pulse sensing (TRPS) and subsequently extracted the EV-RNA. Particles recovered from the two cell lines showed an equivalent size distribution, in the stretched window of an NP250 nanopore (150-900 nm), with no detected significant variation in the mean diameter. At the same time, the relative abundance of secreted particles resulted on average nearly 3-fold enriched (P value 0.011) in the absence of APE1 (Fig. 1A). By contrast, the electrophoretic profile of EV-RNA samples, spanning ∼30-4000 nt in length with major peaks around 150-200 nt (typically indicating the small/fragmented vesicular RNA species (34)), showed a significant reduction of the relative EV-RNA content in the absence of APE1 (∼75%, P value 0.023), when normalized to the corresponding number of particles (Fig. 1B). This variation observed at the level of EV-RNA exacerbated a trend already detectable at the corresponding intracellular level, although to a remarkable lesser extent (∼25% reduction). To better understand the significance of this regulation, we extended the same experiments using other cellular models and profiled the EV-RNA from the A549, human lung adenocarcinoma cell line, transiently transfected for 72 hr with control (SCR) or APE1-targeting siRNAs (siAPE1). As observed in the case of the lymphoma cell lines, the relative RNA abundance decreased by ∼35% in the recovered EV-RNA from APE1-knocked down cells, confirming the positive correlation with APE1 protein levels with an average fluctuation of ∼6% of the EV-RNA content in the SCR condition compared to non-transfected cells (NTC) (Supplementary Fig. S1 and S2). Moreover, we used HeLa cell clones already characterized for APE1 knockdown (HeLa CL3), obtained through the inducible expression of a shRNA targeting the endogenous APE1 (Fig. 1C) (12). As expected from our previous report (22), the APE1 protein is secreted along with EVs and its levels resulted barely detectable in the cell clone compared to the vesicular intraluminal syntenin marker (Fig. 1D). We followed the same pipeline for EV-RNA profiling in these cells and consistently found that the CL3 clone enriched the medium with a 4-fold particle concentration that, in the absence of APE1, lacked about 50% of total RNA compared to SCR cells (Fig. 1E). A reconstitution of the APE1 levels with an shRNA-resistant cDNA sequence partially recovered the amount of EV-RNA content (+15%). This partial rescue was not observed in the case of reconstituted clones expressing mutant APE1K4pleA or APE1NΔ33 encoding plasmids (Supplementary Fig. S3 and S4). These data indicate that APE1 has a remarkable function in determining the intracellular RNA steady-state levels as well as vesicular RNA sorting, possibly as a result of RNA maturation/transport/stability/EV-packaging processes.

**Figure 1:**
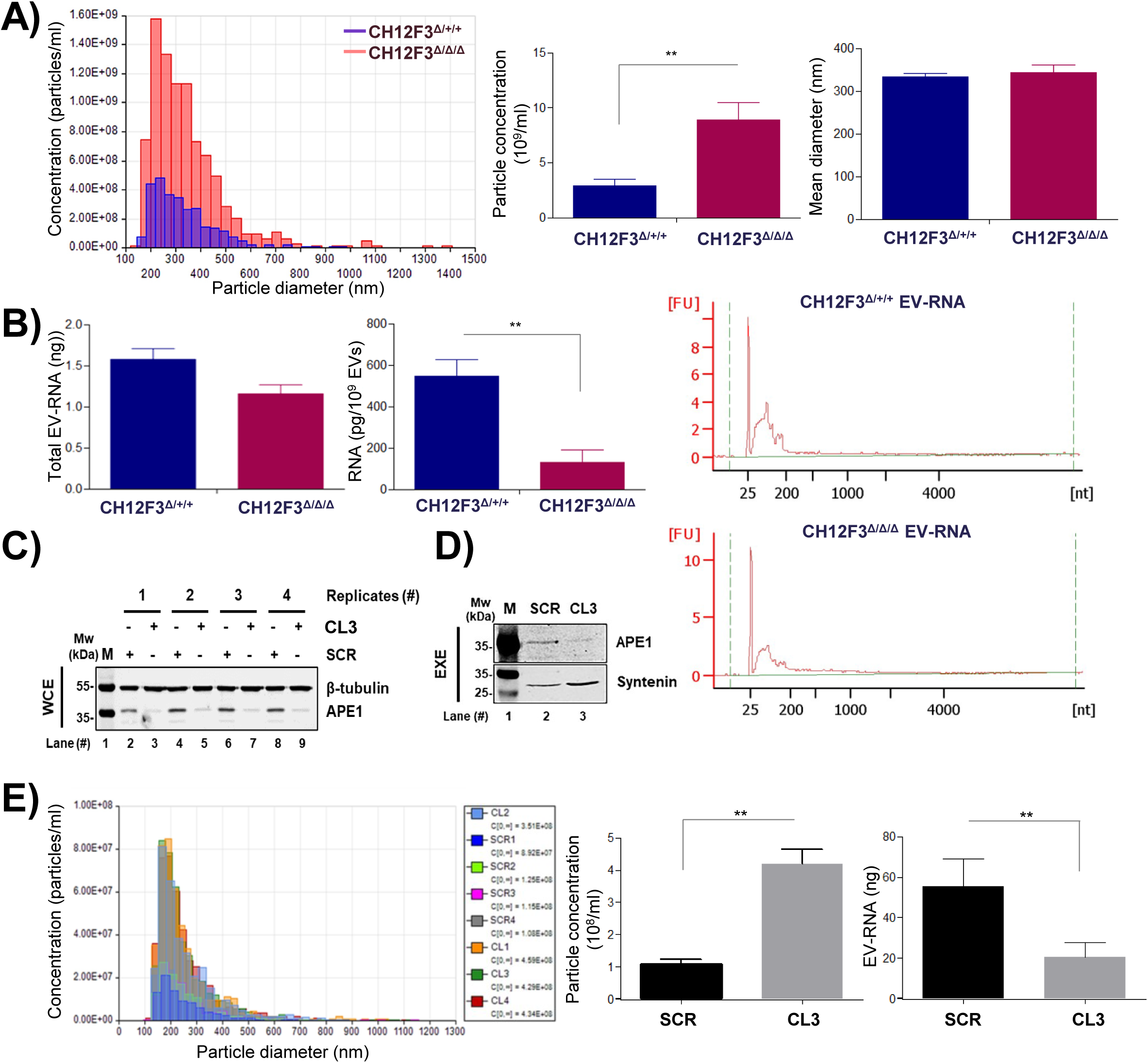
Characterisation of EVs and EV-RNA from APE1-silenced cells. **A**) Representative plot of tunable resistive pulse sensing (TRPS) measurements of EVs recovered from CH12F3Δ/+/+ and CH12F3Δ/Δ/Δ cells. Particles were profiled using qNANO instrument (iZON Science) with an NP250 nanopore (stretched between 44 and 46 mm) calibrated with CPC200 beads (210 nm mean diameter; iZON Science). Samples were diluted 1:2 in PBS 1X before loading and recording the data for 2 minutes in a linear particle count rate. Plots of particle concentration and mean diameter indicate mean and standard deviation resulting from two independent experiments. B) Representative profiles, obtained with Bioanalyzer RNA 6000 Pico Kit (Agilent Technologies), of cellular and vesicular RNA samples extracted from CH12F3Δ/+/+ and CH12F3Δ/Δ/Δ cells and their secreted EVs, respectively. Plots of total and normalized EV-RNA include mean and standard deviation resulting from two independent experiments. C) Western blot analysis for APE1 detection (predicted molecular weight 37 kDa), performed in HeLa inducible APE1 knockdown, indicated as CL3, WCE and its respective control indicated as SCR. β-tubulin (predicted molecular weight 55 kDa) detection was carried out as loading control. The figure shows four experimental replicates. D, western blot analysis for APE1 detection (predicted molecular weight 37 kDa), performed in HeLa inducible APE1 knockdown (indicated as CL3) extracellular vesicles protein extracts (EXE) and its respective control indicated as SCR. Syntenin (apparent molecular weight 32 kDa) detection was assayed as an exosome positive marker. E) TRPS measurements of EVs recovered from SCR and CL3 cells. Biophysical profile and normalized EV-RNA abundance of individual samples are collectively shown. *t-test* was applied to all the comparisons and results were considered statistically significant when *P* value was <0.05 (*), <0.01 (**), <0.001 (***).

### Identification of vesicular miRNAs and ncRNAs affected by APE1-cellular depletion

To investigate the secreted small RNA species regulated by APE1, we performed a small RNA-seq analysis of cell- and EV-RNA (Supplementary Figures S5, and S6) samples deriving from the HeLa clones having stable APE1 reduction (Fig. 1C and D). A principal component analysis (PCA) stratified four well-separated clusters corresponding to HeLa CL3, their parental control cells and the corresponding EV counterparts, with a major variance between EVs and cells (PC1:91%) rather than between cells (PC2: 5%) (Figure 2A), suggesting an active selection of transcripts that are secreted. The distribution of the differentially expressed features (DEFs) is highlighted in the Venn diagram (Figure 2B) as well as in the Volcano Plots (Supplementary Fig. S7), showing the common/APE1-specific transcripts identified in the different conditions. Supplementary Table 1 shows in detail the total number of DEFs, miRNAs and ncRNAs for each performed analysis. Specifically, we identified 257 DEFs (142 up-regulated, 115 down-regulated) in the comparison between Cell CL3 and its control, Cell SCR. Among these, a total of 143 DEFs (112 up-regulated, 31 down-regulated) represent the APE1-silencing-specific expression profile. We then focused on the EV-miR composition, comparing EV CL3 to EV SCR: we identified 325 DEFs (164 up-regulated, 161 down-regulated) with 158 of them (43 up-regulated, 115 down-regulated) being EV-exclusive. Afterward, we investigated the horizontal effect of silencing, comparing EV CL3 to Cell CL3. A total of 3399 DEFs (1152 up-regulated, 2247 down-regulated) were identified, with 568 DEFs (175 up-regulated, 393 down-regulated) being specific to this condition. Lastly, we compared the two controls (EV SCR and Cell SCR) identifying a total of 3989 DEFs (1603 up-regulated, 2386 down-regulated), with 1188 of them exclusive (682 up-regulated, 506 down-regulated). The majority of DEFs are shared between these last two comparisons (EV CL3 vs Cell CL3 and EV SCR vs Cell SCR) and represent the baseline differences existing between the EVs and intracellular transcriptomes. In the control comparison (EV SCR vs Cell SCR) and in the silencing comparison (EV CL3 vs Cell CL3), the percentage of up-regulated miRNAs is on average 10%, while the down-regulated is 15% of the total number of analysed DEFs. Moreover, ncRNAs are present in similar percentages being 15% in up-regulation and 16% in down-regulation. Therefore, the proportions of these two RNA populations in the two conditions remain stable. For what concerns the EV comparison (EV CL3 vs EV SCR), an increase of down-regulated miRNAs is observed up to 42% of the total number of DEFs with a coinciding decrease of ncRNAs (9.9%). Similarly, the same tendency can be observed in the down-regulated portion of Cell comparison (Cell CL3 vs Cell SCR) with 73.9% of DEFs being miRNAs and 3.5% being ncRNAs. However, the up-regulated portion of Cell comparison simultaneously undergoes an increase in the presence of miRNAs (23.9%). These data suggest that APE1 sustains a selection of transcripts that are secreted irrespective of their sub-cellular abundance.

**Figure 2:**
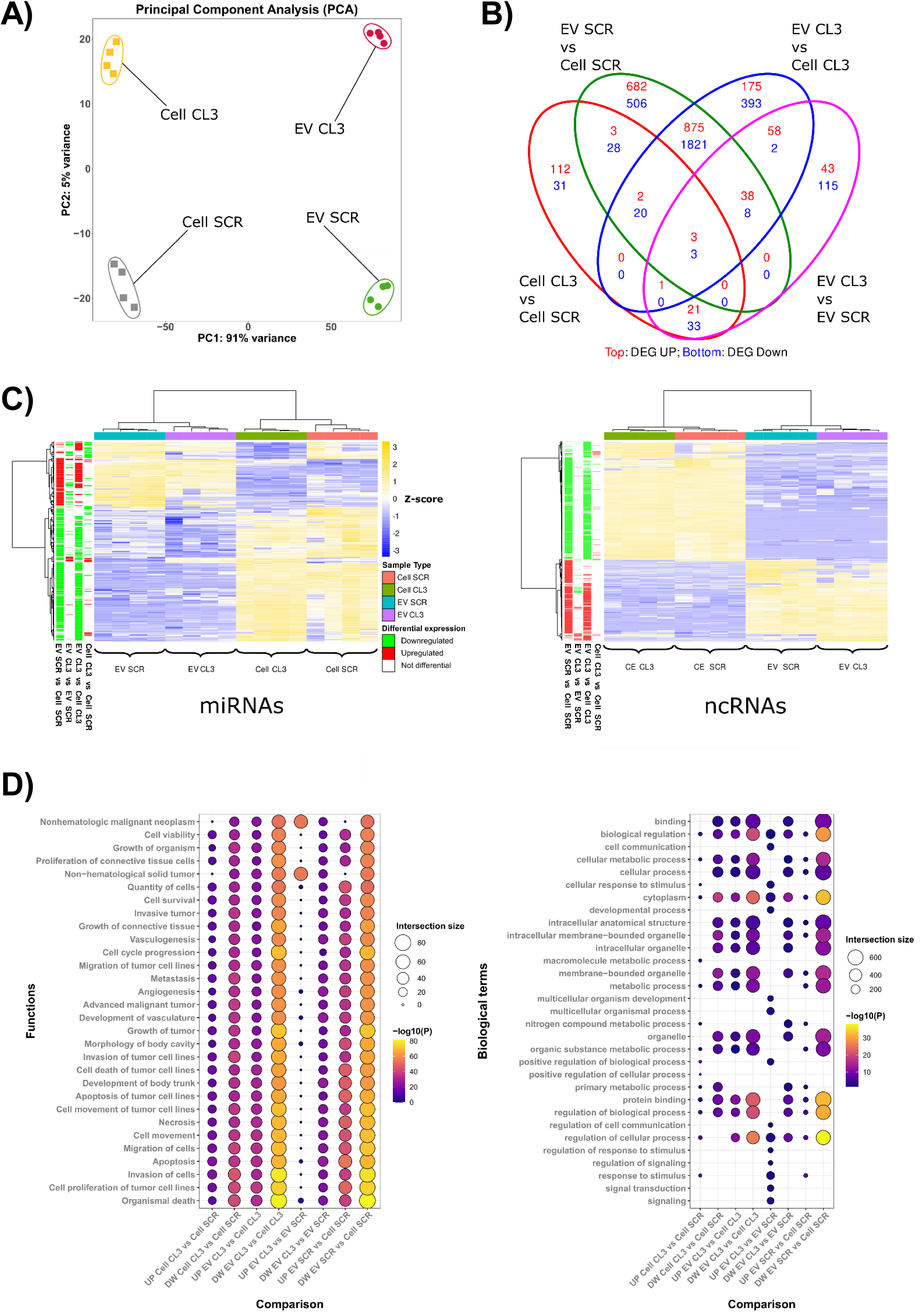
RNA-seq data analyses. A) Principal Component Analysis (PCA) of the expression of the features counts (log2 vst-method normalized counts) for the 12 samples, showing the first versus the second Principal Component (PC). PC1 shows 91% variance and PC2 5%. Each dot represents an EV sample, each square is a cellular sample. Every condition groups together with the same type of samples and indeed it can be observed that the cellular condition is separated from the EVs, as well as the CL3 from SCR. B) Venn of differentially expressed features (DEFs), the numbers in each circle represent the number of DEFs between the different comparisons while the ones overlapping are for mutual DEFs (EV SCR vs Cell SCR is in green, EV CL3 vs Cell SCR is in blue, Cell CL3 vs Cell SCR is in red, EV CL3 vs EV SCR is in purple). The numbers in red represent up-regulated DEFs, blue numbers are relative to the down-regulated portion. C) Heatmap of Z-score of differential miRNAs (left) and ncRNAs (right). “Average” was used as the clustering method and “correlation” for the clustering distance of both rows and columns. Two main clusters of features were obtained: the first comprises the features up-regulated through the samples cluster and the second one comprises the features down-regulated. The track on the right shows differential gene expression between different comparisons (green: down-regulated DEFs; red: up-regulated DEFs; white: no changes in DEFs). D) Balloon plots of enriched molecular functions identified with QIAGEN Pathway Analysis (QIAGEN IPA, left panel) and enriched biological terms characterized with g:Profiler.

The effect of APE1 silencing on differentially expressed ncRNAs and, specifically, on miRNAs can be appreciated through unsupervised clustering (Figure 2C) that separates cell-derived RNAs from EV-derived ones. We further investigated the two subsets, highlighting how miRNAs share a specific expression pattern of up- and down-regulation in EVs, as well as cell-derived, samples (Figure 2C, left). Correspondingly, we observed a similarly shared expression pattern for ncRNAs (Figure 2C, right) between the two sample classes. Both heatmaps highlight specific expression modules associated with APE1 silencing.

In order to understand the possible biological impact of miRNA modulation, we then performed a functional enrichment analysis of the experimentally verified targets obtained from TarBase (36), miRecords (37), and Ingenuity Expert Findings (see Supplementary Tables 2 and 3) (QIAGEN Inc., https://www.qiagenbioinformatics.com/products/ingenuitypathway-analysis). In Figure 2D, the enriched molecular functions identified by QIAGEN Ingenuity Pathway Analysis (QIAGEN IPA, Figure 2D, left panel) and the enriched biological terms resulting from g:Profiler (38) (Figure 2D, right panel) can be seen for each comparison. Interestingly, even if the terms are very similar across comparisons, in all contrasts the downregulated ones are always heightened, underlying the overall impact of APE1 down-regulation. Coherently with the nature of the examined cell models, down-regulated miRNAs in conditions of APE1 silencing are associated with enriched terms involved in cellular growth, cell cycle, migration, and invasion. These data parallel our previous findings about the APE1 prognostic power in tumor progression also through the regulation of microRNA processing mechanisms (12, 17).

Afterwards, explorative motif analysis performed on differentially expressed miRNAs with the MEME suite, suggested the presence of significantly enriched motifs (i.e. GGGGCAGAGA and TCCCAGCC) that could be putative binding sites for the G-quadruplex binding protein APE1, thus providing a mechanism behind the positive correlation between APE1 levels and EV-RNA loading (see Supplementary Table 4, Supplementary Table 5 and Supplementary data). Then we proceeded to investigate EXO-motifs within our DE-miRNAs as these short sequences in miRNAs are recognized mediators for EV-RNA loading.

### EXO-motifs in DE-miRNAs deriving from differential gene expression analysis

Short EXO-motifs sequences are cis-elements known to be specifically recognized by some hnRNPs, such as hnRNPA2B1 and SYNCRIP, and are associated with the sorting of specific miRNAs in EVs (24, 32, 39). On these premises, we investigated the presence of canonical coreEXO-motifs in the 68 down-regulated mature DE-miRNAs (EVDW) upon APE1 silencing in the EV comparison, deriving from the differential gene expression analysis. We retrieved from miRBase all human hairpin sequences and identified a total of 62 unique precursors corresponding to our miRNAs of interest. In order to be unbiased, we searched all possible four-nucleotide short sequences (n=256) both in the precursors and in a list of not differentially expressed miRNAs as background (EVBG, miRNAs n=318, unique precursors n=265). We observed that the EVDW exhibits a statistically significant difference in terms of motif counts with respect to EVBG (Wilcoxon test: W= 1395, p-value < 2.2e-16) showing that EVDW represents a peculiar/distinct population of miRNAs. We decided to further investigate this difference focusing on a single motif level and calculating the ratio of each one between EVDW and EVBG, obtaining 21 motifs with a ratio greater than 1 and p-value≤0.05 using motifcounter tool (40) (Supplementary Table 6). In order to seek the bona-fide EXO-motifs among those, we focused our attention on the subset describing validated coreEXO-motif (n=11) deriving from (41) and thus we found that 6 known coreEXO-motifs were present in our enriched list (AGGG, CCCC, CCUC, GAGG, GGAG, GGCC). Moreover, we further analysed the data, selecting the down-regulated mature DE-miRNAs upon APE1 silencing in EV comparison, that was also not differentially expressed in cellular comparison (EVspecificDW, n=36, number of harpins=34). Also in this case, EVspecificDW exhibited a statistically significant difference in terms of motif counts with respect to the background sequences EVBG (Wilcoxon test: W = 70.5, p-value < 2.2e-16). Interestingly, almost all EVspecificDW harpins (33 out of 34) possessed at least one enriched coreEXOmotifs sequence (Figure 3A). When examining the 6 coreEXO-motifs, we observed that GAGG, GGAG and GGCC aligned with the coreEXO motifs bound by hnRNPA2B1 and that CCUC is the complementary sequence of GGAG (24, 42).

**Figure 3:**
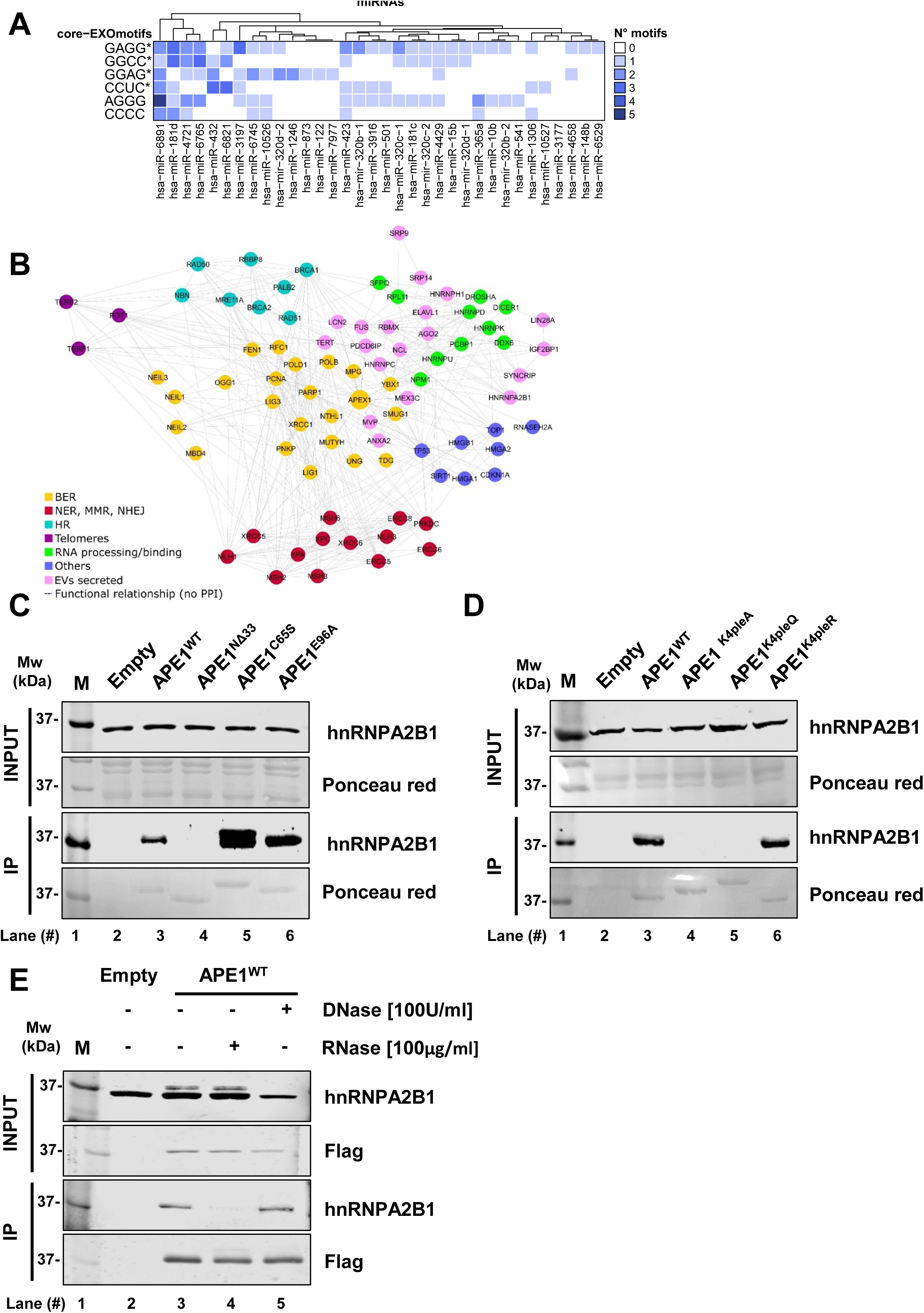
Characterisation of APE1-hnRNPA2B1 interaction. A, Heatmap of enriched coreEXO-motifs bound by down-regulated EV-specific DE-miRNAs subset. Enriched coreEXOmotifs counts are displayed for the EVexclusiveDW hairpin sequences. CoreEXO motifs (GAGG, GGAG and GGCC) bound by hnRNPA2B1 are highlighted by asterisk. Moreover, CCUC is the complementary sequence of GGAG. Colour defines the number of motif occurrences for each hairpin, spanning from white (0 motifs) to black (5 motifs). B, Protein– protein interaction network of proteins involved in DNA repair/stability and in RNA processing. Data, previously published in (2) and now also including RNA-binding proteins identified in EVs and belonging to the APE1 interactome, shows the direct (solid lines) and indirect (dashed lines) interactions existing between BER (yellow nodes), nucleotide excision repair/DNA mismatch repair/non-homologous end joining (NER, MMR, NHEJ, red nodes), homologous recombination (HR, cyan nodes), telomeres (purple nodes), RNA processing/binding (green nodes) and EVs secreted (pink nodes) proteins, as well as a few others (blue nodes) with heterogeneous molecular functions. C, Coimmunoprecipitation (CoIP) analysis performed on JHH-6 cells transfected with APE1WT FLAG-tagged protein and APE1NΔ33, APE1C65S and APE1E96A FLAG-tagged mutant forms. In parallel, cells transfected with PCMV empty vector were analysed as negative control, indicated as Empty. Total cell extracts were immuno-precipitated with FLAG antibody and western blotting analysis was carried out to assay the interaction among APE1 and hnRNPA2B1. Western blot analysis for hnRNPA2B1 (apparent molecular weight 37 kDa) was performed on total cell extracts indicated as INPUT (*upper panel*) and on immunoprecipitated material indicated as IP (*lower panel*). Red Pounceau staining was performed for Input and IP samples as loading controls. D, Coimmunoprecipitation (CoIP) analysis performed on JHH-6 cells transfected with APE1WT FLAG-tagged protein and APE1K4PleA, APE1 K4PleQ and APE1 K4PleR FLAG-tagged mutant forms. In parallel, cells transfected with PCMV empty vector were analysed as negative control, indicated as Empty. Western blot analysis for hnRNPA2B1 (apparent molecular weight 37 kDa) was performed on total cell extracts indicated as INPUT (*upper panel*) and on immunoprecipitated material indicated as IP (*lower panel*). Red Pounceau staining was performed for Input and IP samples as loading controls. E, Coimmunoprecipitation (CoIP) analysis performed on JHH-6 cells transfected with APE1WT FLAG-tagged protein after RNase [100μg/ml] and DNase [100 U/ml] treatments. In parallel, cells transfected with PCMV empty vector were analysed as negative control, indicated as Empty. Total cell extracts were immunoprecipitated with FLAG antibody and western blotting analysis was carried out to assay the interaction among APE1 and hnRNPA2B1. Western blot analysis for hnRNPA2B1 (apparent molecular weight 37 kDa) was performed on total cell extracts indicated as INPUT (*upper panels*) and on immunoprecipitated material indicated as IP (*lower panels*). Western blot analysis using anti Flag antibody was performed for Input and IP samples as loading controls.

In conclusion, we observed putative coreEXO-motif sequences in 58 out of 62 EVDW precursors and 33 out of 34 EVspecificDW precursors, thus supporting once more the hypothesis that APE1 may be directly involved in the EV-miRNA trafficking.

### The APE1 interactome includes several proteins involved in vesicle-mediated transport

In our recent paper (13), the analysis of the APE1 interactome (APE1-PPI) highlighted that APE1 functions can be modulated by different protein interactors. Based on this premise, we analyzed whether APE1-PPI included proteins involved in extracellular vesicle trafficking. We queried the Gene Ontology database through the AmiGO 2 (43–45) (http://amigo.geneontology.org/amigo/landing) and the BioCyc (https://biocyc.org/) websites, defining an exhaustive dataset made of 2208 non-redundant human proteins associated with the “vesicle-mediated transport” term (GO:0016192). A direct comparison with the APE1-PPI interactome allowed us to finally identify hits (n=104) involved in the process (Supplementary Table 7 and Figure 3B). In particular, APE1 interactors were hnRNPA2B1 and NPM1, well-known cargo proteins responsible for miRNA sorting in extracellular vesicles (24, 30, 32). Given this evidence, we, deepened the interplay between APE1 and hnRNPA2B1 to substantiate the mechanism of miRNA recognition and commitment to vesicular RNA secretion.

### Characterization of the APE1-hnRNPA2B1 interaction

Following the results of EV-RNA sequencing, the description of miRNA targets, and the identification of their consensus binding sites, we used our previous knowledge of the APE1 interactome to characterize the interaction between APE1 and hnRNPA2B1. hnRNPA2B1 is a ribonucleoprotein that exerts different functions such as participating in splicing events (46) and RNA decay (47), but, more interestingly, it is involved in regulating specific miRNAs-processing (48) and -sorting into EVs (24). The formation of a protein complex between APE1 and hnRNPA2B1 was already observed in our recent paper (13). Here, we first characterized the APE1 functional domain involved in this interaction through co-immunoprecipitation (CoIP) analyses. CoIP was carried out using the APE1WT Flag-tagged protein and its respective functional mutant forms. First of all, to understand if the interaction was dependent on APE1 redox or nuclease activities or on its ability to interact with protein and RNA molecules, we used: i) the APE1C65S redox mutant, characterized by the substitution of the nucleophilic Cys65 with serine, which abolishes the redox function of the protein (49); the APE1E96A mutant, where the substitution of the Glu96 with alanine compromises its endoribonuclease activity (50), and iii) the APE1NΔ33 deletion mutant, characterized by the deletion of the first 33 aminoacids at the N-terminal unstructured region of the protein, responsible for APE1 interaction with RNA and protein partners (8, 9). Data shown in Fig. 3C demonstrate that APE1 33-N-terminal region is essential for a stable physical interaction between APE1 and hnRNPA2B1. On the contrary, this interaction was not significantly affected in the case of the other two functional mutants, while a serine substitution at position 65 could slightly influence the dynamics of the interaction.

In the N-terminal region of APE1, four Lys residues may undergo *in vivo* acetylation. These post-translational modifications (PTMs) modulate the interaction of APE1 with its protein partners (51, 52). To investigate whether the positive charges (acetylate Lys residues) of the APE1 N-terminal sequence might be essential for the formation of the complex, CoIP analysis in JHH-6 cells was performed, using the mutants APE1K4pleA (52), where the replacement of the Lys27/31/32/35 by Ala residues neutralizes the positive charges of the side chains, and APE1K4pleQ (52), where Lys were substituted with the non-charged Gln residues, mimicking constitutive acetylated residues. Data demonstrated that the positive charges in the N-terminal region are important for a stable interaction (Fig. 3D).

Moreover, since the positive charges of Lys residues are susceptible to acetylation (52), we investigated whether this PTM could affect the interaction between the two proteins. Specifically, we performed CoIP testing on the mutant APE1K4pleR, where the replacement of Lys with Arg residues mimics a non-acetylatable protein form having the opposite biochemical features of the mutant APE1K4pleA (52, 53). As shown in Fig. 3D, we confirmed that the presence of positively charged residues at position 27/31/32/35 is essential for a stable APE1-hnRNPA2B1 interaction.

Finally, to check if this interaction could be mediated by nucleic acids, treatment of the anti-Flag CoIP material with RNAse or DNAse was performed. Results indicate that the APE1-hnRNPA2B1 interaction requires the presence of RNA molecules to stabilize the complex, which is not affected by the presence of DNA, as demonstrated by the sensitivity of the complex formation to RNase-treatment and the resistance of the complex to DNAse-treatment (Fig. 3E). These data demonstrated that, similarly to what previously shown for the APE1-NPM1 complex (53), the APE1-hnRNPA2B1 interaction: i) is mediated by the APE1 unstructured domain, ii) requires charged Lysine residues 27/31/32/35, and iii) is stabilized by the presence of RNA molecules, supporting the hypothesis that APE1 takes part in a dynamic protein complex with hnRNPA2B1 playing a role in miRNA sorting in EVs.

We then validated APE1-hnRNPA2B1 interplay in the regulation of miRNAs containing an EXO-motif (see Supplementary Figure S8 and Supplementary Tables 8 and 9). Comparing all the obtained results, we identified hsa-miR-1246 as: i) the only DEmiRNA, in all the cell lines we tested so far (12, 17), having EXO-motif occurrences in both the precursor and mature forms; ii) a possible interactor of RBPs, based on structural studies showing the presence of EXO-motifs in its sequence (54) and, iii) a highly-expressed ncRNA in EVs derived from many human cancer cell lines (55).

Moreover, upon understanding the importance of hsa-miR-1246, we recovered its validated targets from the TarBase v8.0 database and examined their expression status in a microarray experiment published by our group (10), in which we characterized the transcriptome associated with the loss of APE1 expression in HeLa cells. We identified twelve up-regulated transcripts (ANKRD17, DSP, TAB3, PPP2R1B, NIT1, RBM12, PPP3CA, PLEKHB2, MARCKS, CD164, NEK7 and TNFAIP8; Supplementary Table 10), as validated targets of miR-1246, whose expression is affected by APE1 depletion.

### APE1-hnRNPA2B1 bind to EXO-motif-containing miRNAs

In order to prove that APE1 is able to bind EXO-motif-containing miRs, we performed EMSA analysis using an artificial RNA containing the EXO-motif bound by hnRNPA2B1 (called EXO-miR), a natural human miR-1246, containing the same motif (called hsa-miR-1246), and a scrambled sequence obtained by changing the position of the bases without affecting the overall composition but removing the EXO-motif (called scrEXO-miR-1246) (Fig. 4A). EMSA results, using either the recombinant purified APE1 or hnRNAPA2B1, clearly demonstrate the ability of both APE1 and hnRNPA2B1 to bind the artificial EXO-miR with similar affinity, as well as miR-1246 (Fig. 4B). Also, it appears that the APE1 N-terminal domain is essential for the binding to miR-1246, since the APE1 deletion mutant APE1NΔ33 is not able to stably interact with the miR-1246 probe *in vitro* (Fig. 4C). Titration assays with increasing amount of APE1 and a fixed amount of hnRNPA2B1 demonstrated that the two proteins possibly preserve the stoichiometry on the same consensus sequences (Fig. 4D). In order to demonstrate the specificity of the binding activity by APE1 on miR-1246, we used the scrEXO-miR-1246 probe. Data obtained (Fig. 4E) clearly demonstrated that the EXO-motif present in miR-1246 is required for a stable binding by APE1.

**Figure 4.**
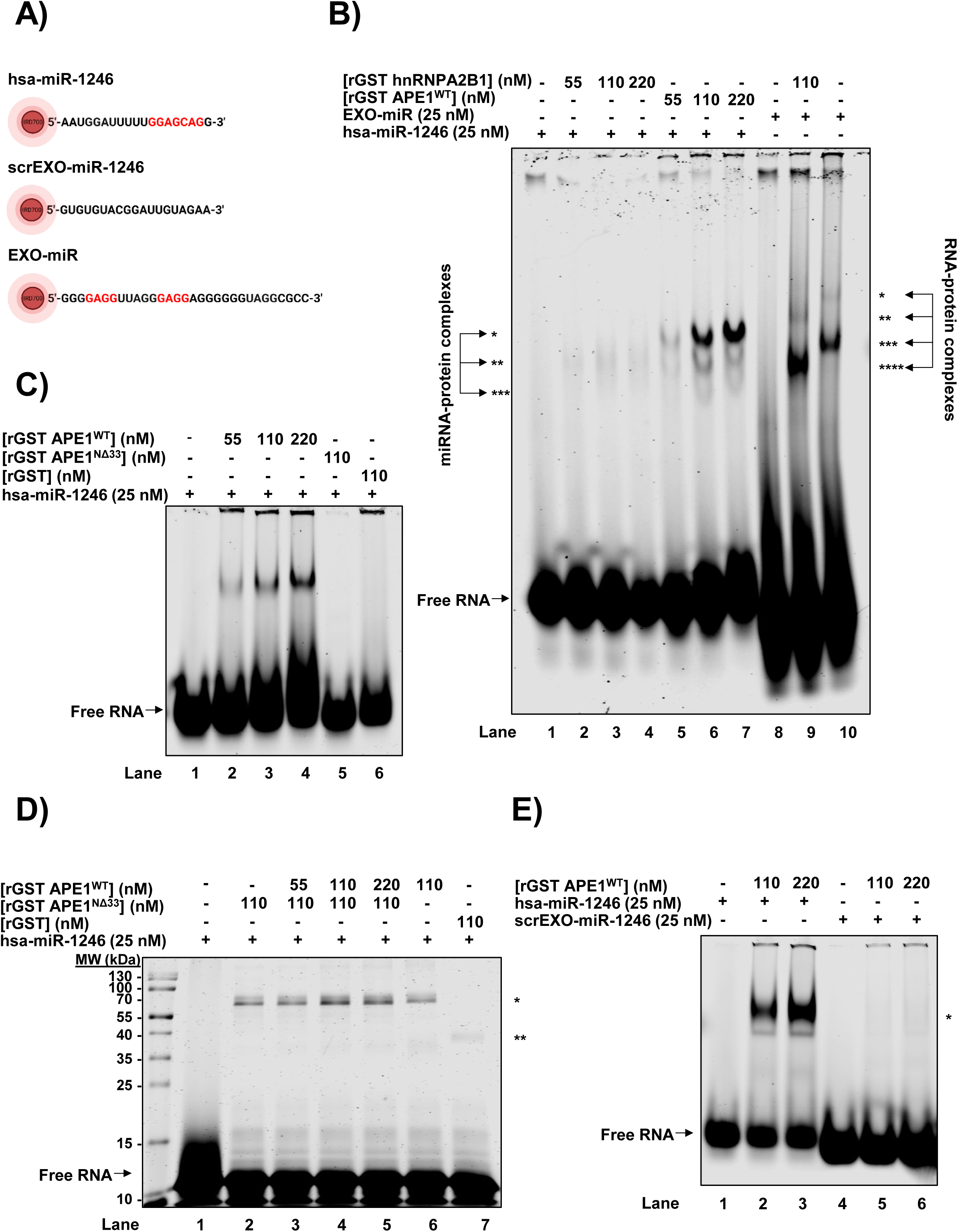
APE1 binds to EXO-motif containing miRNAs. A, EXO-miR and hsa-miR-1246 RNA probes sequence. Schematic representation of IRDye 700 fluorophore labelled ssRNA oligonucleotides called EXO-miR probe (5’-GGGGAGGUUAGGGAGGAGGGGGGUAGGCGCC-3’), hsa-miR-1246 probe (5’-AAUGGAUUUUUGGAGCAGG-3’) and scrEXO-miR-1246 probe (5’-GUGUGUACGGAUUGUAGAA-3’). The 5′ and 3′ ends of each RNA strand are indicated in the figure. Known EXO motif (GAGG) is highlighted in red within the sequence. B, APE1 and hnRNPA2B1 bind to miR-1246 probe *in vitro*. Representative native EMSA 4% polyacrylamide gel of recombinant rGST APE1WT and rGST hnRNPA2B1 proteins. The EXO-miR RNA probe contains two hnRNPA2B1 binding sites and was used as positive control. *, **, *** and **** indicate the signals derived from RNA-protein complexes that display different migration. C, Representative EMSA showing that APE1 deletion mutant APE1NΔ33 doesn’t bind to hsa-miR-1246 probe *in vitro*. Recombinant GST is included as a negative control. * indicates the signal derived from hsa-miR-1246-rGST APE1WT RNA-protein complex. D, Recombinant APE1 and hnRNPA2B1 don’t form ternary complex with hsa-miR-1246 probe *in vitro*. UV-crosslinking followed by SDS-PAGE 15% polyacrylamide gel of recombinant proteins rGST APE1WT, rGST hnRNPA2B1 and rGST APE1NΔ33. Binding to hsa-miR-1246 probe is shown by shift bands (indicated with * and **). E, Representative native EMSA showing that rGST APE1WT is no more able to bind to hsa-miR-1246 if the order of the nucleotide sequence is modified, as in the sequence of scrEXO-miR-1246 probe. * indicates the signal derived from rGST APE1WT-miR-1246 complex.

### miR-1246 features APE1-mediated loading into vesicles

To better characterize the physical association between APE1 and miR-1246, mirroring the sorting process, we performed RNA immunoprecipitation (RIP) experiments. For this purpose, HeLa cells expressing the FLAG-tagged APE1WT protein were analyzed (Fig.5A). Figures 5B and 5C show that both the mature and the precursor form (pri-miR) of miR-1246 co-immuno-precipitated with APE1, proving their physical interaction, compared to HeLa CTR cells, indicated as Empty. Moreover, in order to check if the binding of miR-1246 with APE1 occurred through its N-terminal region and to verify the relevance of the acetylable Lys residues described above on the interaction, RIP analyses were performed also in HeLa cells expressing the mutant forms, APE1K4pleA and APE1NΔ33. Data obtained show that APE1 interacts with miR-1246 and its pri-miR through the N-terminal region of the protein and that substitution of the lysine residues with Ala impairs these interactions. The unstructured N-terminal domain with the acetylable lysine residues 27/31/32/35 seems to have a relevant part in the mechanism.

**Figure 5:**
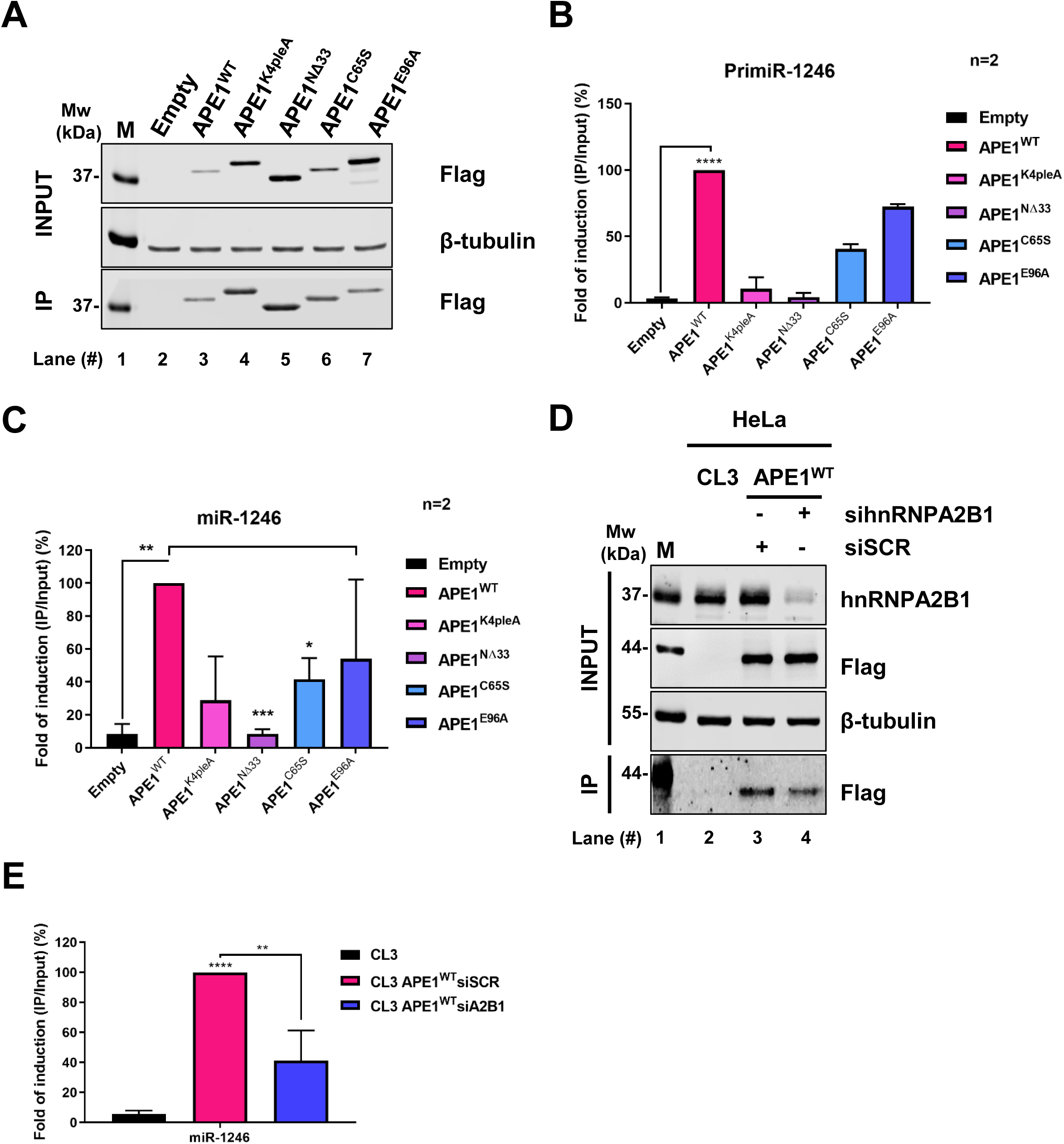
APE1 interacts with miR-1246 cooperating with hnRNPA2B1. A, RNA immunoprecipitation analysis (RIP) performed on HeLa cells transfected with APE1WT FLAG-tagged protein and APE1K4PleA, APE1NΔ33, APE1C65S, APE1E96A FLAG-tagged mutant forms. In parallel, cells transfected with PCMV empty vector were analysed as negative control. RIP was carried out using FLAG antibody. Western blot analysis using anti-Flag antibody was performed for Input (*upper panel)* and IP (*lower panel*) samples and western blot for β-tubulin was performed in INPUT samples (*middle panel*) as loading control. B, qPCR analysis for Pri-miR-1246 was performed on Input and IP RNAs derived from RIP experiment. The histogram shows the fold of induction of IP/Input expressed in percentage. Data are expressed as mean ± SD of two independent replicas, ****p < 0.0001. C, qPCR analysis for miR-1246 was performed on Input and IP RNAs derived from RIP experiment. Histogram shows the fold of induction of the ratio IP/Input expressed in percentage. Data are expressed as mean ± SD of two independent replicas, *p < 0.05, **p < 0.01, ***p < 0.001. D, RNA immunoprecipitation analysis was performed on inducible APE1 knock down HeLa cells overexpressing APE1WT FLAG-tagged protein (indicated as CL3 APE1WT), transiently silenced for hnRNPA2B1 or for a siSCR as a control. In parallel, inducible APE1 knock down HeLa cells (indicated as CL3) were used as RIP negative control. The immunoprecipitation was performed using FLAG antibody. Western blot analysis was carried out in protein samples derived from RIP experiment. The detection of hnRNPA2B1 (apparent molecular weight 37 kDa) and β-tubulin (apparent molecular weight 55 kDa) as loading control were performed in Input samples. The detection of APE1WT-FLAG protein was performed using anti FLAG antibody in IP samples (*lower panel*) and in Input samples as control. The molecular mass (MW) expressed in kDa is shown on the left side of the image. E, qPCR analysis for miR-1246 was carried out in Input and IP RNAs derived from RIP experiment. Input and IP RNAs derived from inducible APE1 knock down HeLa cells were used as RIP negative control. Data are expressed as a fold of induction of the ratio IP/Input. Data are expressed as mean ± SD, **p < 0.01, ****p < 0.0001.

Afterward, in order to evaluate the role of hnRNPA2B1 in the interaction between miR-1246 and APE1, we performed RIP in reconstituted HeLa APE1WT cells upon transient depletion of hnRNPA2B1 through siRNA downregulation (Fig.5D, and E). As shown in Figure 5E, the silencing of hnRNPA2B1 impaired the interaction between APE1 and miR-1246, demonstrating that hnRNPA2B1 is essential for this interplay.

We then checked whether the exosomal sorting of miR-1246 was dependent on the APE1 protein together with its protein partner hnRNPA2B1. For this purpose, we analysed cellular (Cell) and exosomal (EXO) miR-1246 expression in HeLa cells transiently depleted of APE1, hnRNPA2B1, or both proteins in combination, using specific siRNAs (Figure 6A) and reporting the EXO/Cell value expressed in percentage, obtained from qRT-PCR analyses of miR-1246. As shown in Figure 6B, APE1 silencing significantly affected the exosomal amount of miR-1246. Likewise, APE1 silencing caused a significant increase of intracellular miR-1246 (Cell miR-1246) with respect to the exosomal counterpart, in agreement with the RNA-seq data. The same trend was also observed using the A549 cell line (Supplementary Figure S9). Higher levels of Cell miR-1246 were also observed in the condition of double knockdown (siAPE1/hnRNPA2B1) (Fig. 6D). The same trend was not observed when APE1 was knocked down in combination with its protein partner NPM1 (Fig. 6C and 6D), underlying the specificity of the event depending on the APE1 interacting partner.

**Figure 6:**
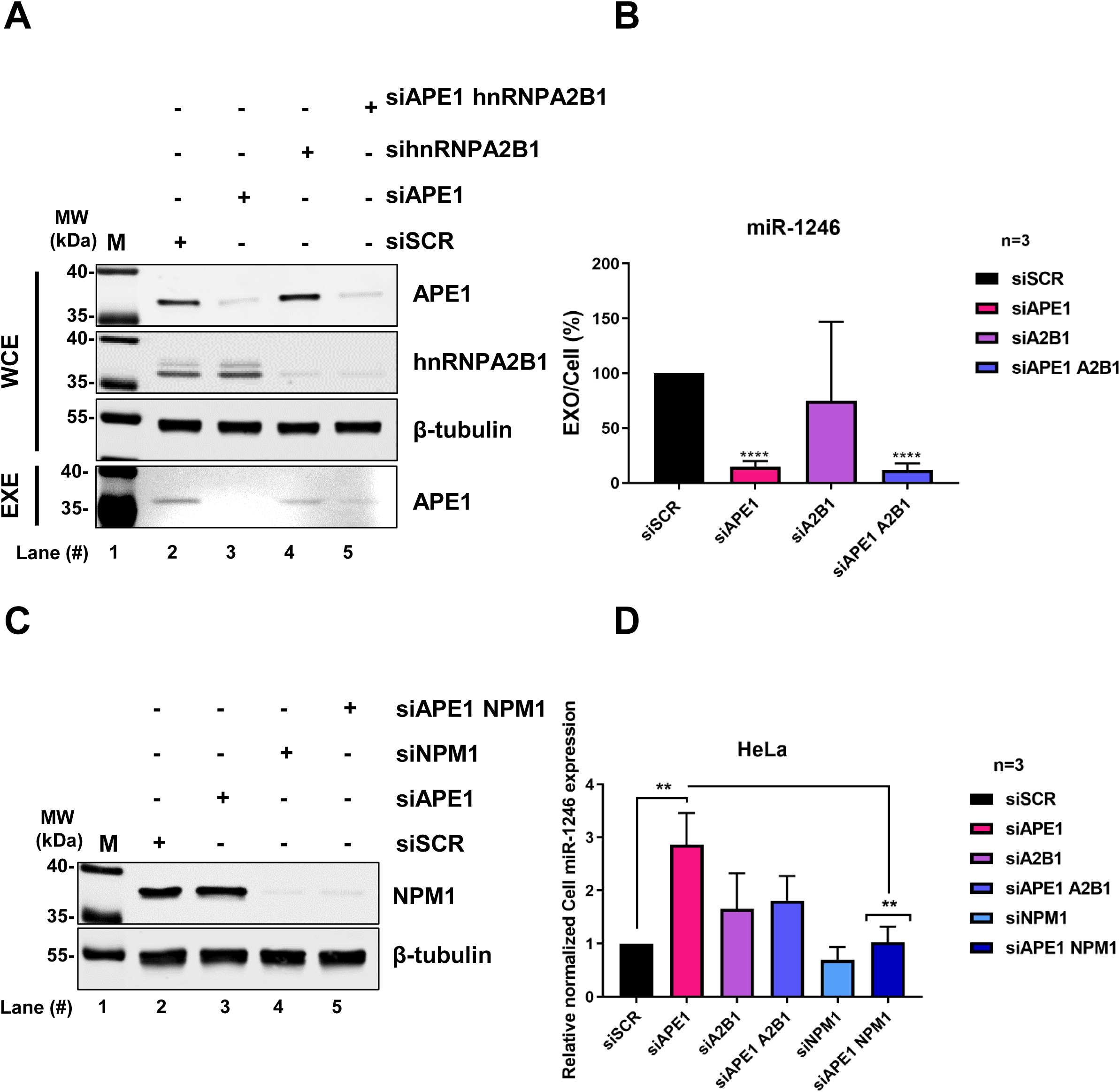
APE1 downregulation dysregulates miR-1246 cellular pool. A, Western blot analysis was carried out on WCE and EXE of HeLa cells silenced for APE1, hnRNPA2B1 or both proteins in combination. The detection of endogenous APE1 (apparent molecular weight 37 kDa) is shown in *upper panel*, follows the detection of hnRNPA2B1 (apparent molecular weight 37 kDa). β-tubulin (apparent molecular weight 55 kDa) detection was carried out as loading control. The detection of exosomal APE1 (apparent molecular weight 37 kDa) is shown in the *lower panel*. The molecular mass (MW) expressed in kDa is shown on the left side of the image. B, qPCR analysis for miR-1246 expression carried out on endogenous (Cell) and exosomal (EXO) miRNAs derived from HeLa cells silenced for APE1, hnRNPA2B1, and both proteins in combination. The silencing with a siRNA scramble (siSCR) was performed as control. Data represent the ratio EXO / Cell miR-1246 expression expressed in percentage. miR-16-5p was used as a reference gene only for endogenous RNA. The exosomal RNA was normalized by Qubit quantification. Data are expressed as mean ± SD of three independent replicas, ****p < 0.0001. C, Western blot analysis carried out on WCE of HeLa cells silenced for APE1, NPM1 and both proteins in combination. The detection of NPM1 (apparent molecular weight 37 kDa) and β-tubulin (apparent molecular weight 55 kDa) as loading control are shown. The molecular mass (MW) expressed in kDa is shown on the left side of the image. D, qPCR analysis for miR-1246 expression carried out on endogenous RNAs (indicated as Cell miR-1246) derived from HeLa cells silenced for APE1, hnRNPA2B1, NPM1 and APE1 in combination with hnRNPA2B1 or with NPM1. The silencing with a siRNA scramble (siSCR) was performed as control. Data represent the relative normalized Cell miR-1246 expression. miR-16-5p was used as a reference gene. Data are expressed as mean ± SD of three independent replicas, **p < 0.01.

Altogether, these data support a model in which the APE1-hnRNPA2B1 interaction plays a central role in sorting specific miRNAs within extracellular vesicles.

To corroborate our data demonstrating that APE1 regulates the miRNA-exosome sorting, thanks to its capability to interact with the EXO-motifs, we validated: a) cellular (Cell) and exosomal (EXO) miR-1246 and miR-181d-5p both bearing the EXO-motifs GGAG (Supplementary Figures 10 A-D) and b) APE1 capability in regulating EV sorting of miRNAs bearing GC-rich consensus motifs, testing one of the eight microRNAs having this feature, i.e. miR-320e (Supplementary Figure 10 E and F).

### TCGA expression analysis of APE1 and hnRNPA2B1 and prognostic significance of identified vesicular microRNAs

Considering that APE1 and hnRNPA2B1 are both negative prognostic factors in several tumors (56, 57), we wanted to evaluate and compare their expression levels in different cancers. For this purpose, gene expression correlation analysis was performed by consulting *The Cancer Genome Atlas* (TCGA) datasets. As shown in Supplementary Figure S11, there is a significant positive correlation between the expression levels of APE1 and hnRNPA2B1 in most of the examined tumors (n=28). The levels of correlation are reported in (Supplementary Table 11). To explore the prognostic power of candidate miRNAs in different cancer types, we first retrieved both miRNA expression and clinical data for different cancers in which APE1 is known to be overexpressed: Lung Adenocarcinoma (TCGA-LUAD) and Cervical Squamous Cell Carcinoma and Endocervical Adenocarcinoma (TCGA-CESC). We then computed, for each dataset, a prognostic index (PI) to stratify subjects into two different classes based on both miRNA expression and miRNA association with overall survival. In particular, the subject’s PI was computed using multivariate Cox regression coefficients and the expression values of selected miRNA for which expression data were available (eight miRNAs respectively in TCGA-LUAD and TCGA-CESC, Fig.7A and Fig. 7C). Subjects were then stratified into two groups, High risk, and Low risk, based on *p-value* optimization. Finally, Kaplan-Meier curves and a log-rank test were used to summarize data and compute the associated *p-value*. The overall survival rates were significantly different between High risk and Low risk groups in the TCGA-LUAD (Fig. 7B, *n* High risk = 148 and *n* Low risk *=* 279*, p-value* = 1* 10-06) and in the TCGA-CESC datasets (Fig. 7D, *n* High risk *=* 155 and *n* Low risk *=* 118*, p-value* = 0.002). Taken together these findings support the potential prognostic value of the miRNA signature in the two cancer types analysed.

**Figure 7:**
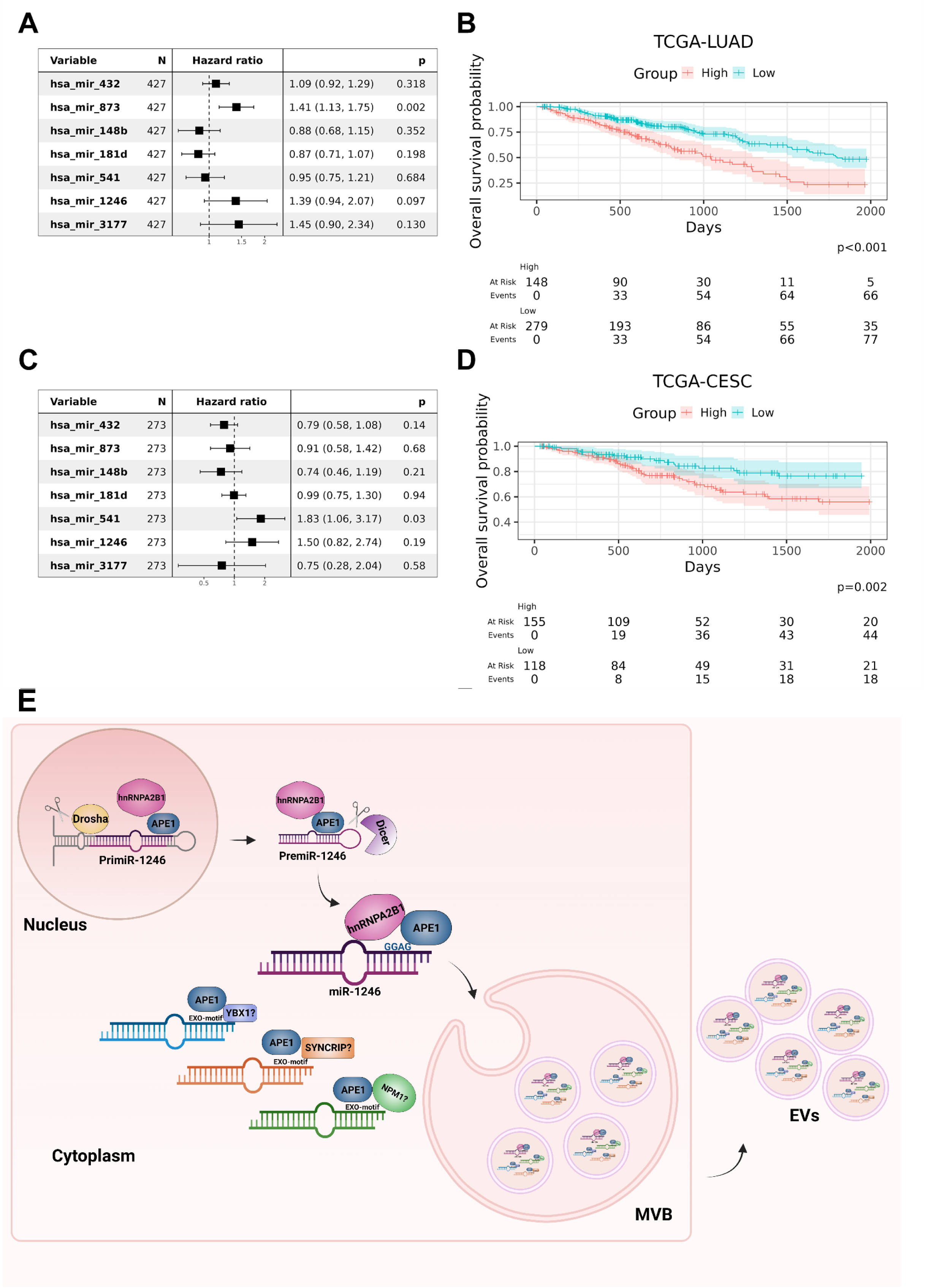
Prognostic significance of identified vesicular microRNAs: Forest plot of the multivariate cox proportional hazard model in the (A) TCGA-LUAD and (C) TCGA-CESC datasets. The *Variable* column contains the miRNA name, the *N* column contains the total number of samples analysed, the *hazard ratio* column contains the distribution of the hazard ratio and the *P* column the p-value associated with each miRNA. Prognostic value of the miRNA signature in (B) TGCA-LUAD and (D) TCGA-CESC datasets. Kaplan–Meier plot showing the different overall survival rates of subjects belonging to the “High risk” and “Low risk” groups, stratified based on the Prognostic Index calculated from the miRNA signature. Associated *p-*values are reported under each plot. E) Representative model of miR-1246 sorting process, mediated by the complex APE1-hnRNPA2B1, starting from the nucleus, where miR-1246 transcription and processing begins, and ending with EVs secretion. miRNAs bearing the EXO-motifs recognised by APE1 and the other APE1 protein interactors, implicated in vesicles-mediated transport such as NPM1, SYNCRIP and YBX1 are also shown.

## DISCUSSION

Understanding the mechanisms responsible for the enrichment of specific RNA subsets into extracellular vesicles represents a crucial missing point that can explain the paracrine function of secreted EVs and non-coding RNAs in physiological and pathological conditions like, for instance, tumor progression. Notably, these mechanisms could be at the basis of chemoresistance processes determining the secretion of a specific EV-RNA pattern, therefore representing druggable axes to potentially interfere with the tumor microenvironment. The identification of pivotal mechanistic factors would guide and dramatically enhance our capacity to non-invasively intercept disease biomarkers, given the emerging role of EVs in modern liquid biopsy. In this view, EVs offer the fascinating opportunity to reach these mechanisms, like the activity of specific intracellular factors determines a selected EV content, showing *per se* a reduced complexity compared to the cellular machinery (58).

In this study, we unveiled the molecular mechanism for sorting motif-enriched miRNAs into EVs involving the DNA-repair APE1 protein. By using different APE1-depleted human cancer cell lines (i.e. HeLa, A549), in which we transiently or inducibly silenced the cellular APE1 protein through specific shRNAs, or using APE1-knock out mouse lymphoid cell lines (CH12F3Δ/+/+ and CH12F3Δ/Δ/Δ cells), we showed that APE1 dosage positively correlates with the intracellular miRNA steady-state as well as with vesicular RNA levels. Further experiments with different APE1-defective mutants clearly showed a central role of the Lys residues in position 27/31/32/35, present in the N-terminal unstructured APE1 domain, for the observed effects. Quantitative miRNA-seq analysis using cellular and vesicular RNA counterparts of APE1-depleted or -expressing cells allowed us to identify specific consensus miRNA sequences affected by the dosage of the APE1 protein. Additional cellular and biochemical approaches, such as RIP- and expression analyses performed in cells or by using purified recombinant proteins, allowed us to identify the EXO-motif sequence of miR-1246 as a representative binding site for APE1, undoubtedly establishing the unexpected, non-canonical role of this DNA repair protein in the EV-miRNA sorting.

We observed the presence of EXO-motifs in miRNAs, following APE1 silencing, in EVs as compared to the control EVs. Interestingly, the EXO-motifs were also observed in miRNA precursors and in particular in the miR-320 family, which is known to be involved in different tumor entities (59), and in the miR-181 family, which are implicated in cancer and neurodegeneration (58). In order to further highlight the role that miRNAs play after the silencing of APE1, we decided to investigate only those exclusive to EVs and not present at the cellular level. What we observed was a higher number of EXO-motifs present in the 34 miRNA precursors, finding at least one statistically significant EXO-motif in 33 of them. These findings imply that APE1 is directly implicated in the process of EV-cargo loading of miRNAs, in particular in those possessing one or more EXO-motifs, whose presence is cumulatively EV-related.

Importantly, we characterized the activity of APE1 in the interplay with hnRNPA2B1. Members of the hnRNP family (i.e. hnRNPA2B1, hnRNPC1, hnRNPG, hnRNPH1, hnRNPK and hnRNPQ) along with some additional RNA-binding proteins (ALIX. AGO2, ANXA2, FUS, LIN28, QK1, IGF2BP1, MEX3C, NCL, YBX1, SRP9/14 and TERT), emerged as pivotal players involved in RNA-packaging events of specific miRNAs within EVs (31). We here show that the APE1 and hnRNPA2B1 proteins physically interact and constitute a regulatory axis for the packaging and secretion of a subset of miRNA-containing motifs. In Figure 7E we propose a representative model of miR-1246 sorting and secretion process mediated by the complex APE1-hnNRPA2B1.

Our deepened analysis of miRNA sequences cannot lead to excluding other RNA species that could be sensitive to the APE1/hnRNPA2B1 dosage. Recently, the hnRNPA2B1 protein has also been implicated in the EV-packaging of motif-enriched long RNA transcripts (60) or the sorting of miRNAs into microvesicles upon interaction with caveolin 1 (61), meaning that the interplay of these proteins can occur in specific phases of the maturation/acquisition of vesicles. In this view, we showed the consistent contribution of APE1 to protein:protein/RNA interaction networks possibly regulating the endosomal compartment and the selection of motif-enriched miRNA species. Dedicated experimental pipelines are needed to systematically analyze the contribution of the APE1/hnRNPA2B1 axis on different RNA species. Taking advantage of all the data produced during this study, we argue that a prevalently nuclear protein like APE1 closely follows bound RNA targets during maturation, transport, stabilization, target recognition and up-to-date EV-packaging. We have also shown that APE1 is part of the EV-proteome and is still functional against oxidized RNAs *in vitro*, supporting the hypothesis that a specific enrichment on GC-rich motifs could require levels of control from protein recruiters that *determine or protect* the final commitment of these non-coding RNAs. Our findings open a new intriguing scenario for the potential role of APE1 in the cell-to-cell communication, possibly intervening on both RNA- and DNA-substrates, and the RNA quality control process for secreted EVs.

As an exhaustive target, we characterized the interaction of APE1 with miR-1246, already explored in the sorting process within EVs (54) and for its paracrine role in tumor progression of different cancer types (55). miR-1246 has been found to be oncogenic in lung, cervical, liver, colorectal, breast, pancreatic and ovarian cancers through mechanisms regulating GSK3β, JAK/STAT, PI3K/AKT, RAF/MEK/ERK, Wnt/β-catenin, THBS2/ MMP and NOTCH2 pathways. Interestingly, APE1 is overexpressed in all the above-mentioned cancer types (2, 13, 62) and in lung and liver cancers increased level of secreted APE1 in affected patients was found (19, 20). These findings support the hypothesis of a relevant role of APE1 in the tumorigenic process through paracrine mechanisms involving specific oncogenic miRNAs. Further work, along these lines, is required to better prove this hypothesis and to extend our observations to other secreted miRNAs. Knowledge of the molecular details responsible for this novel function of APE1 in the sorting process of specific onco-miR could help in designing new anti-cancer strategies based on targeting the specific interaction rather than targeting the general redox-/repair-functions of APE1, as it has been done to now (63–66). These perspectives, in our opinion, will give the possibility to better translate APE1 functions from benchtop to bedside with the aim of developing new personalized medicine approaches. We acknowledge that our results represent a preliminary hypothesis in this sense, which should be experimentally validated through additional *in vivo* studies. Our data suggest that modulating the expression levels of APE1 may affect miRNA sorting within EVs and, therefore, we postulate associations with clinical responses to anticancer drug treatments. In parallel, we provide a first rationale to potentially test APE1-regulated miRNAs as novel prognostic biomarkers in multiple cancers, possibly with concomitant combinations with new APE1 inhibitors. Further exploration of the recognized associations is expected to improve drug effectiveness and to identify interesting therapeutical combinations for precision medicine.

## MATERIAL AND METHODS

### Cell culture

Cells used for this publication are cultured as reported in (12, 22)

### Whole cell extracts preparation (WCE)

WCE were obtained as indicated in (22)

### Extracellular vesicles isolation

We systematically used a nickel-based isolation procedure for EV isolation, according to previous literature (34, 35) and, specifically, for secreted APE1 (22). Media were supplemented with 10% FBS exosome-depleted (Thermo Fisher Scientific), 100 U/ml penicillin, 100 μg/ml streptomycin (Euroclone). Polydisperse particles were then characterized by tunable resistive pulse sensing (TRPS) with a qNANO Instrument (iZON Science) as already reported (31, 34), and using an NP250 nanopore stretched between 44 and 46 mm calibrated with CPC200 beads (210 nm mean diameter; iZON Science). Samples were diluted 1:2 in PBS 1X before loading and recording the data for 2 minutes in a linear particle count rate. We also confirmed the reproducibility of the results obtained by RNAseq, validating by qPCR not only RNA samples derived from EVs extracted with NBI, but also, as an alternative procedure, using RNA samples derived from particles extracted with Exosome Isolation kit (Thermo Fisher Scientific), according to the manufacturer’s instructions. In this latter case, exosomes were isolated from medium supplemented with 10% FBS exosome-depleted (Thermo Fisher Scientific), 100 U/ml penicillin, 100 μg/ml streptomycin (Euroclone) to strengthen the validation of our results. In the case of western blotting, EV protein extracts were prepared as reported in (22).

### EV-RNA isolation, library preparation, and RNA sequencing

Vesicular RNA was extracted by mixing EVs with TRIZOL reagent (Merk), then adding 20 μl of chloroform and incubation for at room temperature for 5 min after shaking. Phases were separated by centrifugation at 12,000*g* for 15 min at 4 °C, and the aqueous phase was recovered to proceed with RNA purification using columns for Single Cell RNA Purification (Norgen). Following DNAse treatment with RNase-Free DNase Set (Qiagen), RNA concentration and purity were assessed by Nanodrop 2000 spectrophotometer (Thermo Scientific™) while its integrity (RNA integrity number, RIN) was assessed by gel capillary electrophoresis using Agilent 2100 Bioanalyzer instrument (Agilent Technologies, CA, USA), following the manufacturer’s instructions.

A total of 5 ng of EV-RNA/sample and 100 ng of cellular RNA/sample were used as a template for the preparation of cDNA libraries using the QIAseq miRNA Library Kit (Qiagen) following the manufacturer’s instructions. Relative sample yields were determined by Qubit with the DNA High Sensitivity kit (Life Technologies). Quantified libraries were mixed at an equimolar ratio and sequenced on the NovaSeq6000 platform (Illumina, USA) using a 100-bp flow-cell. RNA-seq was performed in quadruplicate (GEO number GSE230874) from HeLa SCR and CL3 clones and derived EVs.

### Statistical analysis

Statistical analyses were performed by using the Student’s t-test in GraphPad Prism software. When P<0.05 data were considered as statistically significant. The Barnard’s Unconditional Test and the Mann-Whitney test were performed using the Barnard (https://CRAN.R-project.org/package=Barnard) and the stats (https://www.R-project.org) packages in the R/Bioconductor environment.

Methods concerning bioinformatics analysis and wet lab experiments are included in the Supplementary Information.

## DATA AVAILABILITY

Data regarding the RNA-sequencing experiment is publicly available at either raw and count level at NCBI Gene Expression Omnibus Database with GEO accession number GSE230874 (the following secure token has been created to allow review of record GSE230874 while it remains in private status: ozghkmgyjxsvjqv).

## ACKNOWLEDGEMENTS

The work was supported by a grant from Associazione Italiana per la Ricerca sul Cancro (AIRC) [grant number IG19862] to G. T. and through the support of the Departmental Strategic Plan (PSD) of the University of Udine—Interdepartmental Project on Artificial Intelligence (2020-25) and by additional grants from the University of Udine (‘Bando Ricerca Collaborativa’ granted by European Community - NextGenerationEU) and from the Consorzio Interuniversitario Biotecnologie - C.I.B. – (ex D.M. MUR n.1059 del 09.08.21: "L’INNOVAZIONE DELLE BIOTECNOLOGIE NELL’ERA DELLA MEDICINA DI PRECISIONE, DEI CAMBIAMENTI CLIMATICI E DELL’ECONOMIA CIRCOLARE”) to G.T.

## AUTHOR CONTRIBUTIONS

G.T. and V. D. A designed and conceived the study and supervised the experiments; G.M. performed most of the experiments, analyzed the data, and critically contributed to the interpretation of the results; M.N. contributed to the experiments on EVs RNA characterization and prepared the NGS libraries, F.F. contributed to the experiments on EV-RNA characterization; G.C., E.D., N.G., S.P. performed the bioinformatics analysis; S.P. supervised the Bioinformatic analyses; V. D. S. performed the NGS sequencing; M.D.G. performed EMSA experiments; G.A. helped with EMSA analysis; G.M., V. D. A., S.P. and G.T. mainly wrote the manuscript. All authors critically read and approved the final version of the manuscript.

## COMPETING INTERESTS

The authors declare no competing interests.

## ADDITIONAL INFORMATION

**Correspondence** and requests for materials should be addressed to Gianluca Tell

## Supporting Information

### Supplementary Data

#### Motifs analysis on differentially expressed miRNAs

Speculating on direct control of APE1 to bind miRNAs based in a sequence-specific manner, we started a motifs discovery analysis by comparing the EV CL3 against EV SCR, focusing separately on up- and down-regulated miRNAs. In order to conduct the analysis with the most unbiased possible approach, we applied different programs belonging to the MEME Suite (MEME, STREME and XSTREME, TOMTOM, MCAST, MAST) (1), as well as Weeder2 (2), to find consensus sequences in differentially expressed miRNAs. The results of all these analyses are summarized in Supplementary Table 4.

Using MEME, we identified a single enriched motif, GGGYDSWGRKGS (meme_motif1, E-value=1.9e-002), present in 56 out of 68 examined down-regulated miRNAs (see Supplementary Table 5). No statistically significant results were found for up-regulated miRNAs. We then applied STREME on the down-regulated miRNAs, using the unchanged miRNAs as control, discovering five statistically significant motifs that, however, we deemed as unfit due to the definition of a consensus just for the first or last three nucleotides (i.e AAAGS). Similar analyses were repeated: i) on downregulated miRNAs with unchanged miRNAs and upregulated ones as control sequences, using the same settings, and then on upregulated miRNAs using as a background either ii) on unchanged miRNAs alone or iii) together with the downregulated ones, identifying for these three analyses five significant motifs each with the same characteristics of the ones found in the first STREME investigation. An XTREME analysis was then performed on both down- and up-regulated miRNAs with either unchanged miRNAs or the up/downregulated ones as a background, outputting what was already observed in the STREME analysis: short motifs with a consensus made of just a couple of nucleotides. Finally, we used Weeder2 both on the up-regulated miRNAs, obtaining no significant motif, and on the downregulated miRNAs, identifying two identity matrices for the following motifs: GGGGCAGAGA (weeder_motif1) and TCCCAGCC (weeder_motif2). In detail, weeder_motif1 was found in 8 sequences: hsa-miR-4429, hsa-miR-423-5p, hsa-miR-10526-3p, hsa-miR-320e, hsa-miR-320d, hsa-miR-320c, hsa-miR-320a-3p and hsa-miR-320b (position p-value < 0.0001), while weeder_motif2 was found in 7 down-regulated miRNAs: hsa-miR-7977, hsa-miR-4429, hsa-miR-320e, hsa-miR-320d, hsa-miR-320c, hsa-miR-320a-3p and hsa-miR-320b (position p-value < 0.0001). Notably, some of them belong to the same family, such as the hsa-miR-320 family (hsa-miR-320a-3p, hsa-miR-320b, hsa-miR-320c, hsa-miR-320d and hsa-miR-320e). In conclusion, our analyses suggest that differentially expressed miRNAs exhibit significantly enriched motifs that could be putative binding sites for the APE1 protein, thus providing a mechanism behind the positive correlation between APE1 levels and EV-RNA loading.

#### APE1 regulated miRNA containing EXO-motif are validated by the APE1-hnRNPA2B1 interplay

Based on previous knowledge (3) about the existence of two conserved, enriched short sequences (EXO-motifs) identified in mature miRNAs that could mediate specific RNA-sorting into exosomes, we searched for the presence of miRNAs containing the two aforementioned EXO-motifs (motif1: BGVS ([GUC]G[ACG][GC]); motif2: BCCD ([GUC]CC[UGA])) (Supplementary Fig. S8) within the datasets of cellular dysregulated miRNAs (DE-miRNAs) in APE1 knocked down cells available in our laboratory. Specifically, we queried DE-miRNAs identified in the following cell lines: JHH-6 (see Supplementary Table 8 and (4)), A549 (5) and HeLa (6). We recovered the mature human miRNA sequences of interest from miRBase (7) (ftp://mirbase.org/pub/mirbase/CURRENT/mature.fa.gz) and then used the FIMO program (http://meme-suite.org/tools/fimo) from the MEME-Suite (8, 9) to identify all the EXO-motif occurrences in the examined datasets. In JHH-6 cells, we found only four DE-miRNAs having one or two copies of each EXO-motif, while in A549 and HeLa cells (both APE1-depleted and H2O2-treated, Supplementary Table 8) we identified tens of DE-miRNAs harboring either EXO-motif1 or 2 in their mature sequences (Supplementary Table 9), 33-50% containing both.

### Supplementary Figures

**Supplementary figure S1).**
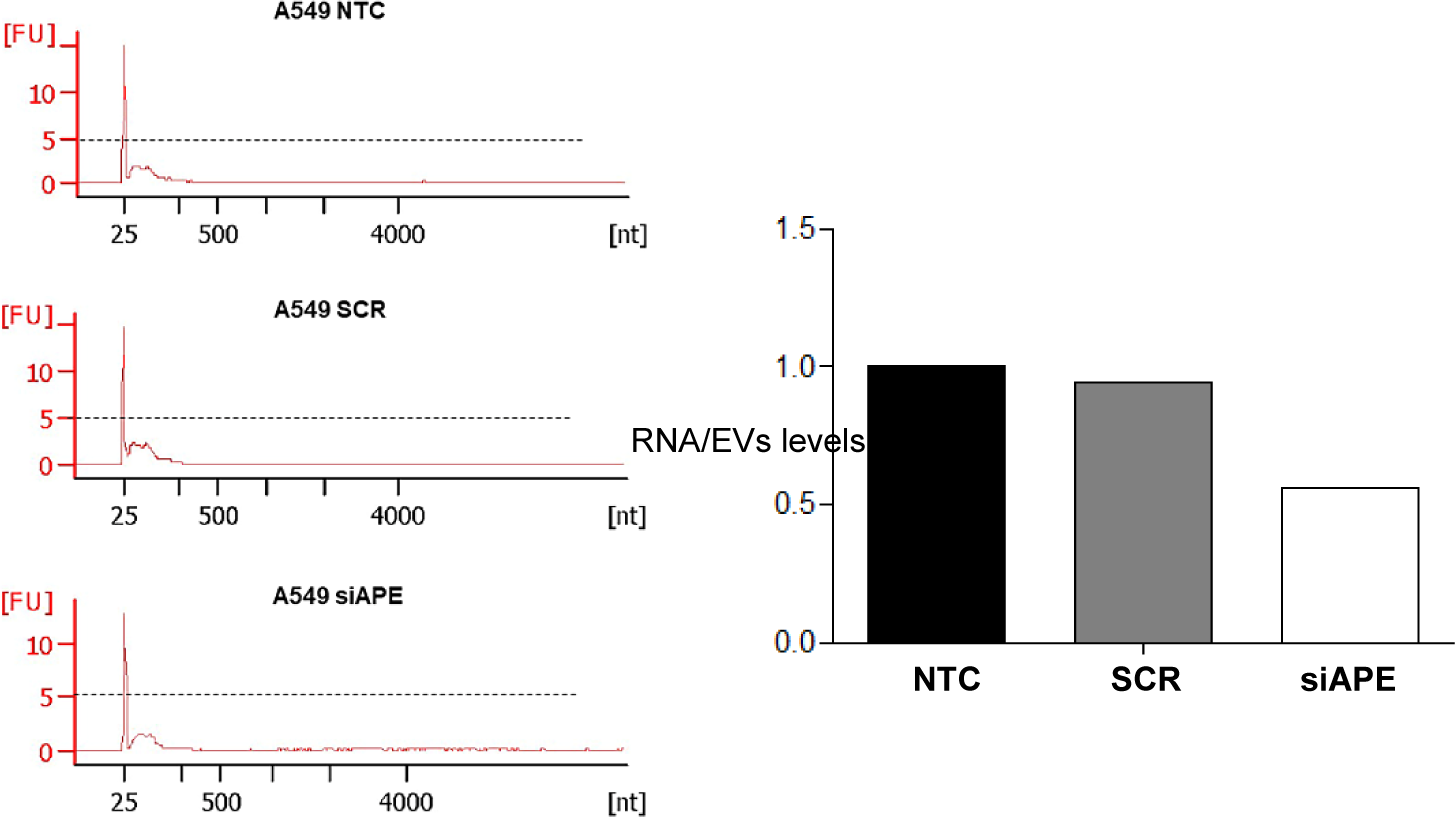
Vesicular RNA profiling recovered from A549 media not transfected cells (NT), A549 siSCR and A549 siAPE1 transfected cells, on the right histograms representing normalized EV-RNA levels.

**Supplementary figure S2).**
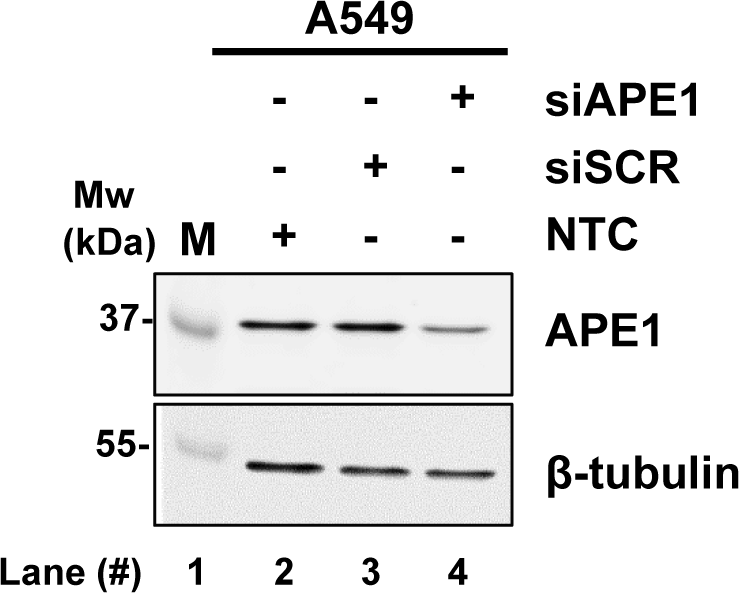
Western blot analysis for APE1 detection performed in A549 silenced for APE1 for 72 hours and the respective SCR control. β-Tubulin detection was carried out as loading control.

**Supplementary figure S3).**
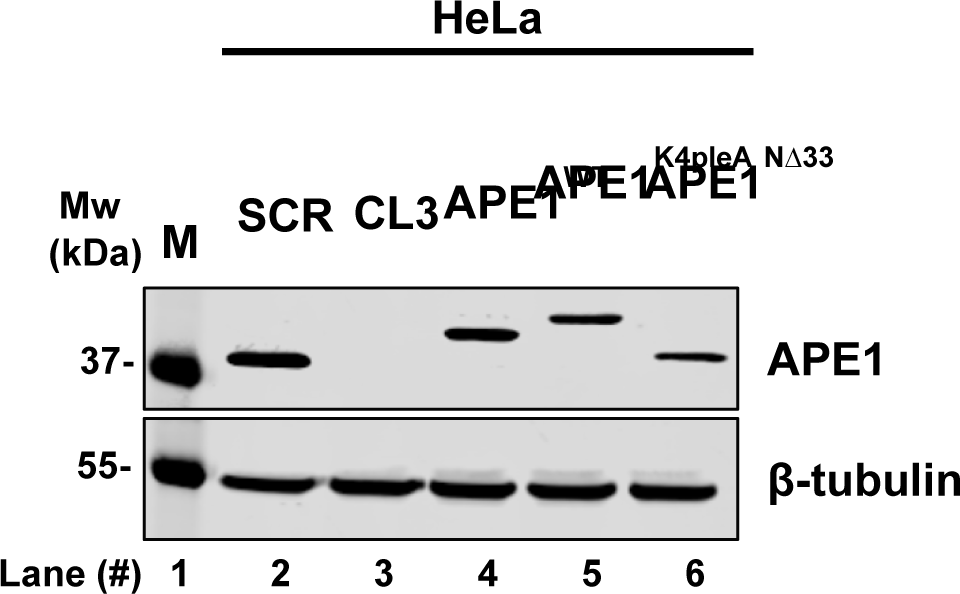
Western blot analysis for APE1 detection was carried out in HeLa inducible knock down cell lines. HeLa SCR indicates cell line which expresses endogenous APE1 protein levels. HeLa CL3 indicates cell line in which an inducible downregulation of endogenous APE1 was carried out. The CL3 reconstituted cell lines with APE1^WT^, APE1^K4pleA^, APE1^NΔ33^ are also shown. Western blot for β-tubulin detection was performed as loading control.

**Supplementary figure S4).**
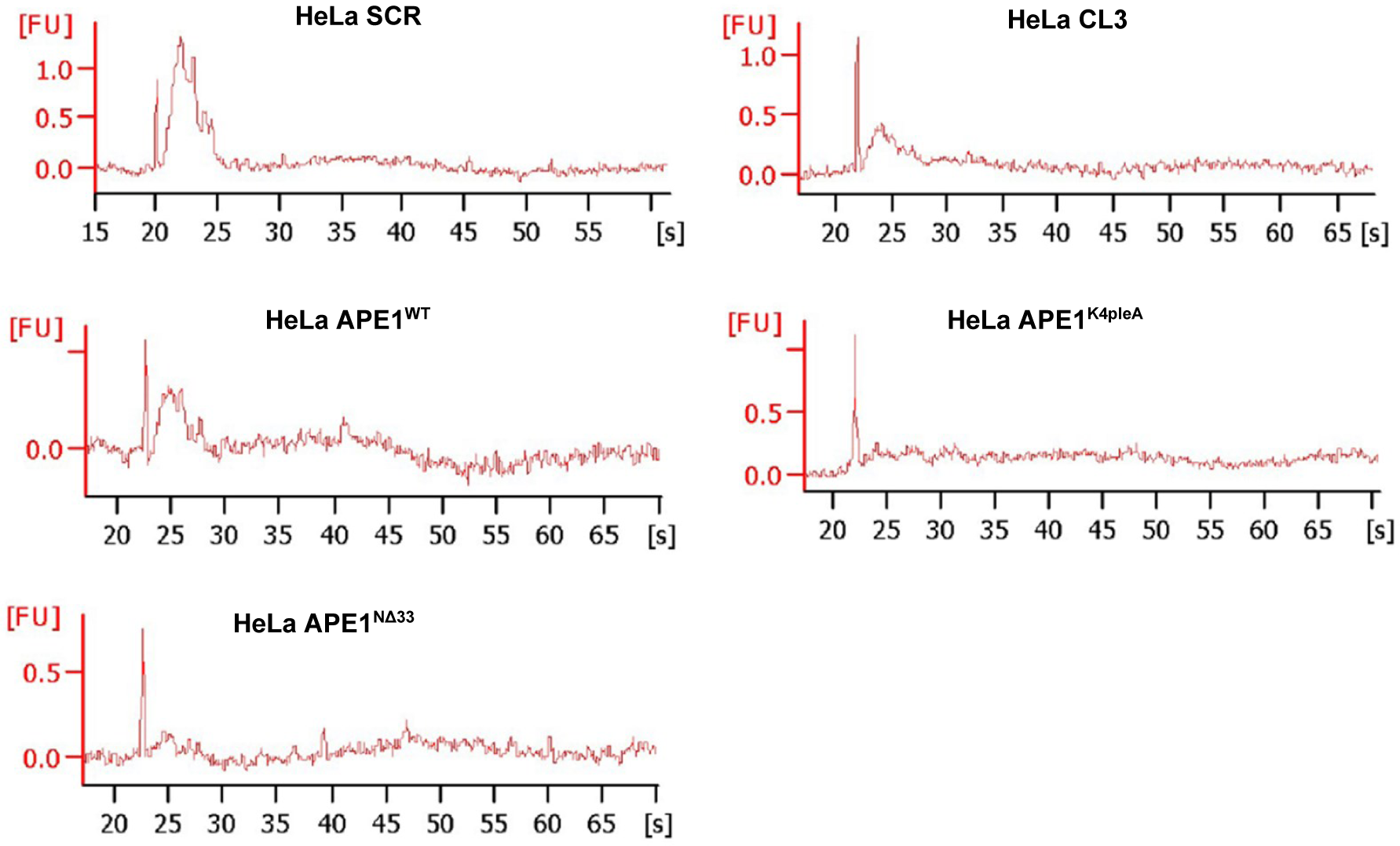
Vesicular RNA profiling recovered from media of HeLa SCR, CL3 and reconstituted APE1 WT, K4pleA and NΔ33 clones.

**Supplementary figure S5).**
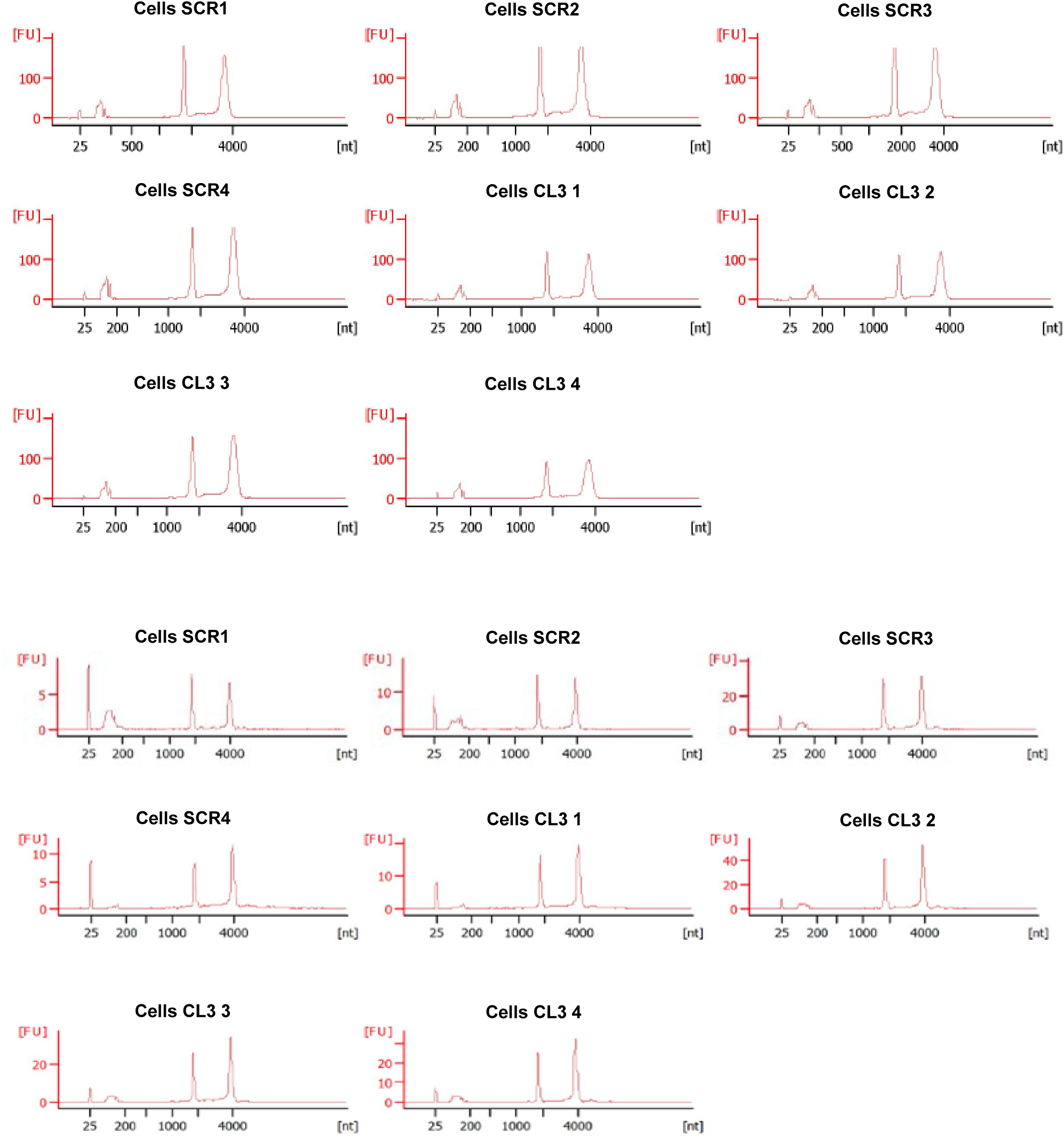
RNA profiles obtained from HeLa SCR, CL3 cells.

**Supplementary figure S6).**
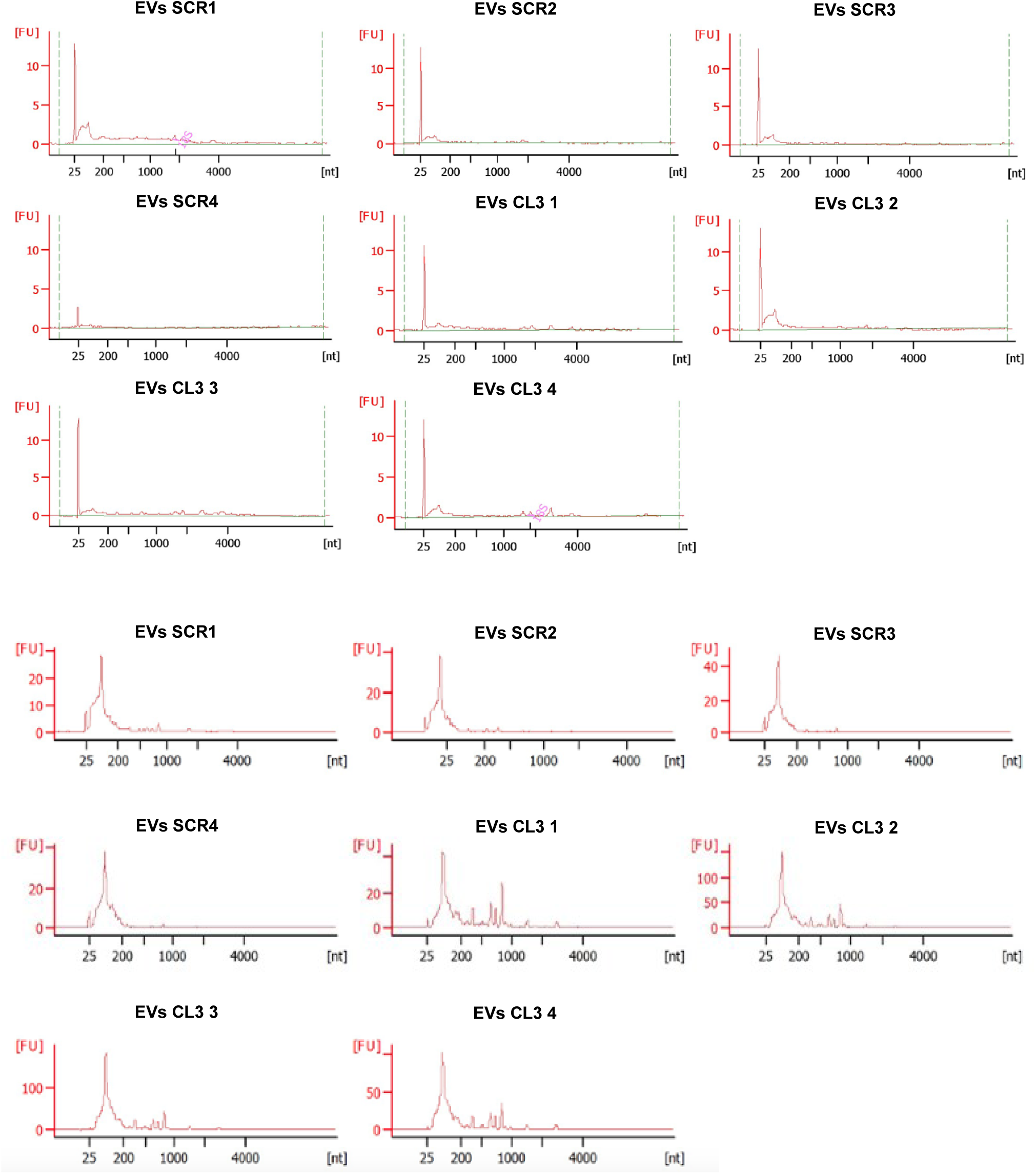
RNA profiles obtained from HeLa SCR, CL3 EVs.

**Supplementary figure S7).**
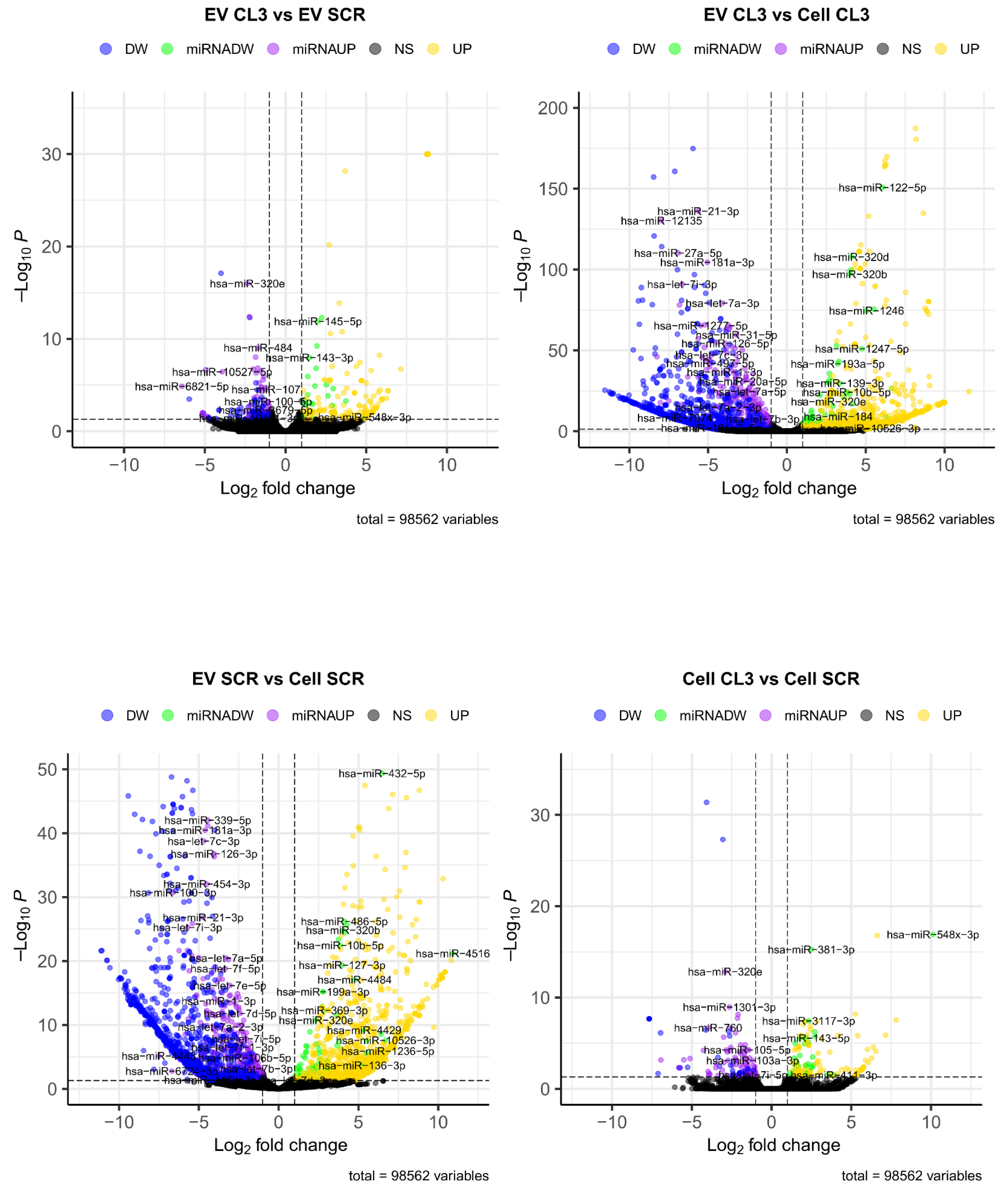
Volcano plots of the distribution of differentially expressed features (DEFs) in the four performed comparisons in the small RNA-seq. Measurement of feature expression log2fold-change is present on the X-axis against a measure of statistical significance (-Log10 (P-value)] on the Y-axis. Differential features are established at |Fold change| ≥ 1 and P-value <0.05 (yellow: up-regulated DEFs; purple: up-regulated miRNAs; blue: down-regulated DEFs; grey: no changes in DEFs; green down-regulated miRNAs).

**Supplementary figure S8).**
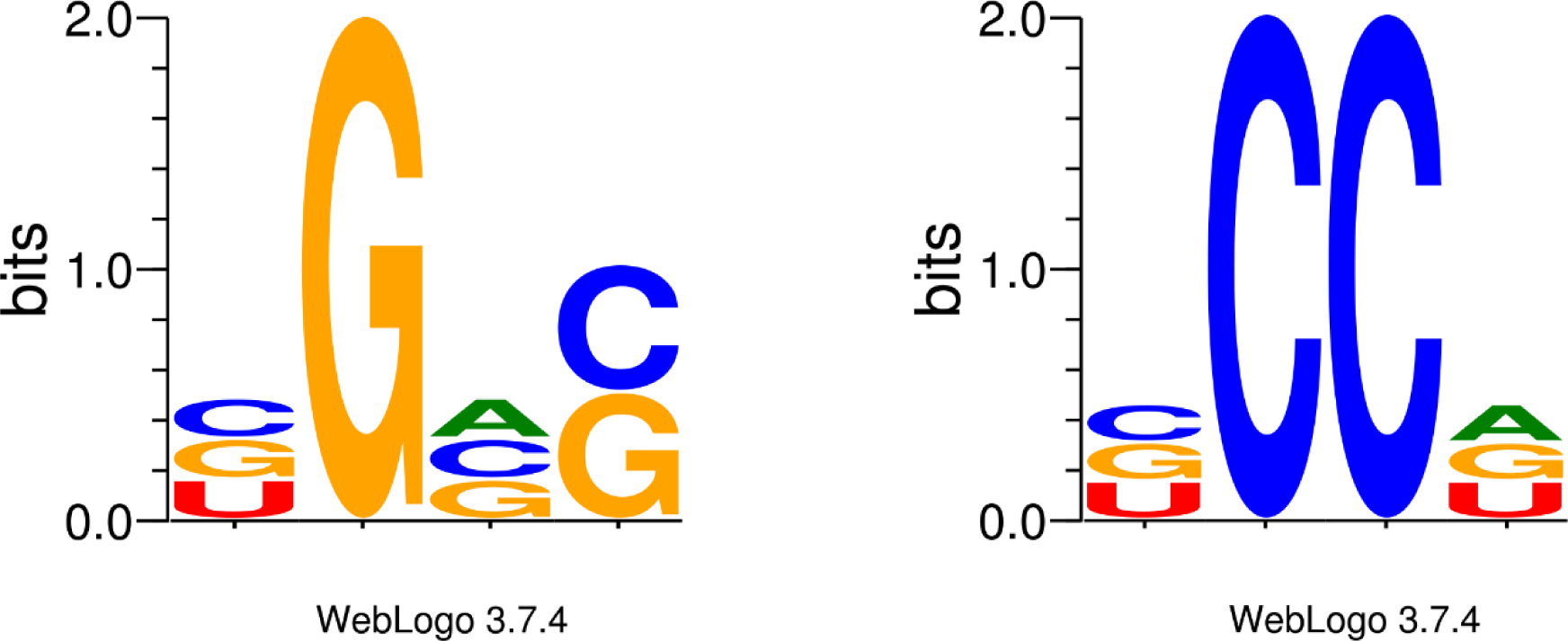
Logo representation of the investigated EXO-motifs 1 *(left)* and EXO-motifs 2 *(right)*.

**Supplementary figure S9).**
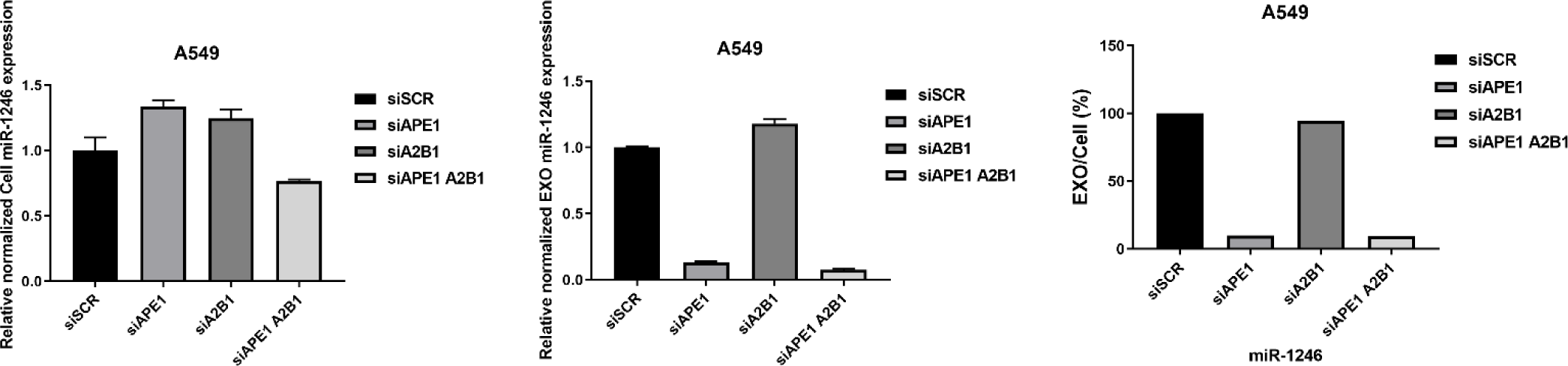
*Left panel*), qPCR analysis for miR-1246 expression carried out on endogenous RNA (indicated as Cell miR-1246) derived from A549 cells silenced for APE1, hnRNPA2B1, and APE1 in combination with hnRNPA2B1. The silencing with a siRNA scramble (siSCR) was performed as control. Data represent the relative normalized Cell miR-1246 expression. miR-16-5p was used as reference gene. *Middle panel)* qPCR analysis for miR-1246 expression carried out on exosomal RNA (indicated as EXO miR-1246) derived from A549 cells silenced for APE1, hnRNPA2B1, and APE1 in combination with hnRNPA2B1. The silencing with a siRNA scramble (siSCR) was performed as control. Data represent the relative normalized EXO miR-1246 expression. *Right panel*), qPCR analysis for miR-1246 expression carried out on endogenous (Cell) and exosomal (EXO) miRNAs derived from A549 cells silenced for APE1, hnRNPA2B1, and both proteins in combination. The silencing with a siRNA scramble (siSCR) was performed as control. Data represent the ratio EXO / Cell miR-1246 expression expressed in percentage. miR-16-5p was used as reference gene only for endogenous RNA. The EXOsomal RNA were normalized by Qubit quantification.

**Supplementary figure S10).**
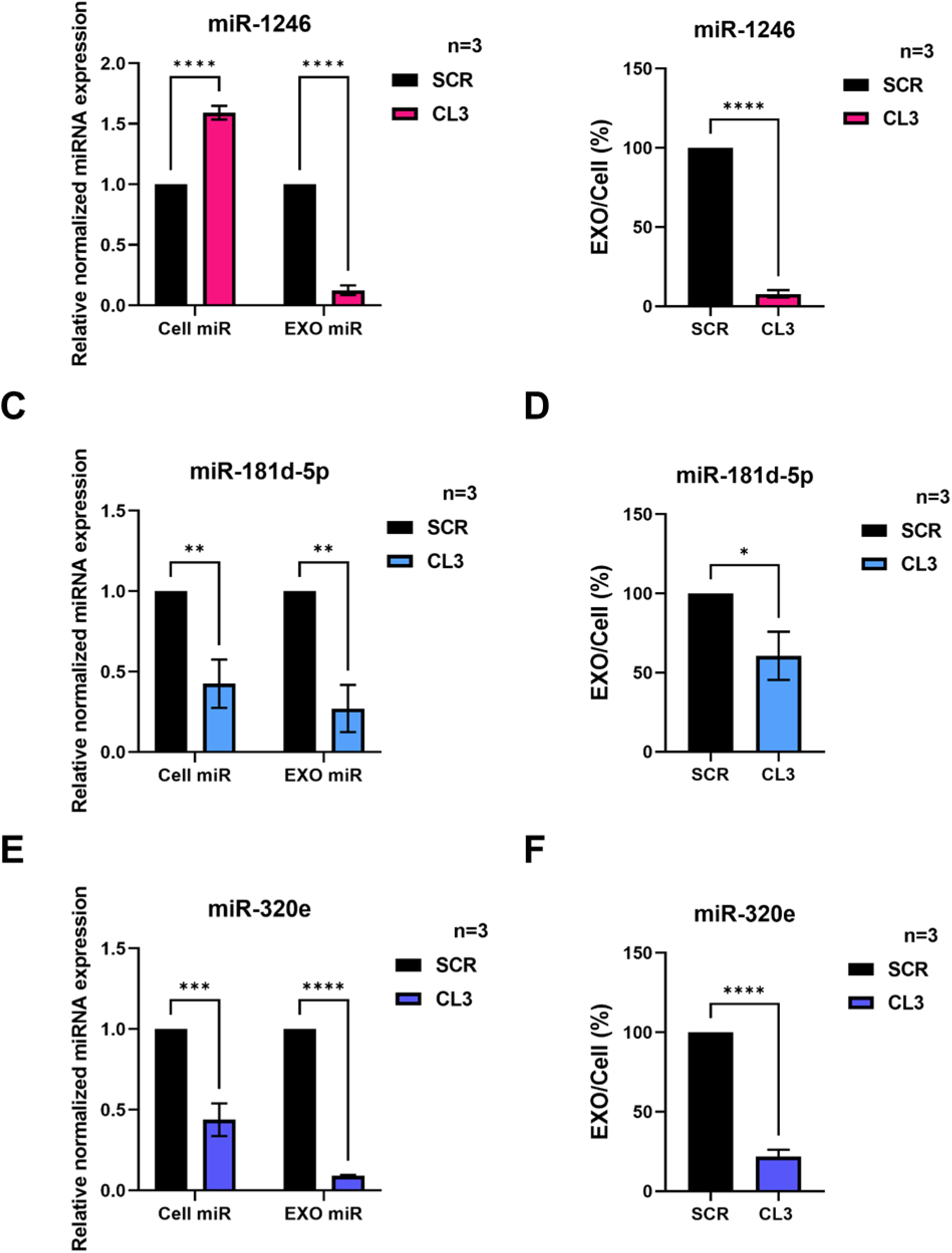
Impairment of miRNAs bearing Exo motifs sorting upon APE1 depletion. A, qPCR analysis for miR-1246 expression carried out on endogenous (Cell) and exosomal (EXO) miRNAs derived from HeLa CL3, and its respective control indicated as SCR. Data are expressed as relative normalized miRNA expression respect to SCR control. miR-16-5p was used as reference gene only for endogenous RNA. The exosomal RNA were normalized by Qubit quantification. Data are expressed as mean ± SD of three independent replicas, ****p < 0.0001. B, histograms representing the ratio EXO / Cell of miR-1246 expression expressed in percentage. Data are expressed as mean ± SD of three independent replicas, ****p < 0.0001. C, qPCR analysis for miR-181d-5p expression carried out on endogenous (Cell) and exosomal (EXO) miRNAs derived from HeLa CL3, and its respective control indicated as SCR. Data are expressed as relative normalized miRNA expression respect to SCR control. miR-16-5p was used as reference gene only for endogenous RNA. The exosomal RNA were normalized by Qubit quantification. Data are expressed as mean ± SD of three independent replicas, **p < 0.01. D, histograms representing the ratio EXO / Cell of miR-181d-5p expression expressed in percentage. Data are expressed as mean ± SD of three independent replicas, *p < 0.05. E, qPCR analysis for miR-320e expression carried out on endogenous (Cell) and exosomal (EXO) miRNAs derived from HeLa CL3, and its respective control indicated as SCR. Data are expressed as relative normalized miRNA expression respect to SCR control. miR-16-5p was used as reference gene only for endogenous RNA. The exosomal RNA were normalized by Qubit quantification. Data are expressed as mean ± SD of three independent replicas, ***p < 0.001; ****p < 0.0001. F, histograms representing the ratio EXO / Cell of miR-320e expression expressed in percentage. Data are expressed as mean ± SD of three independent replicas, ****p < 0.0001.

**Supplementary figure S11).**
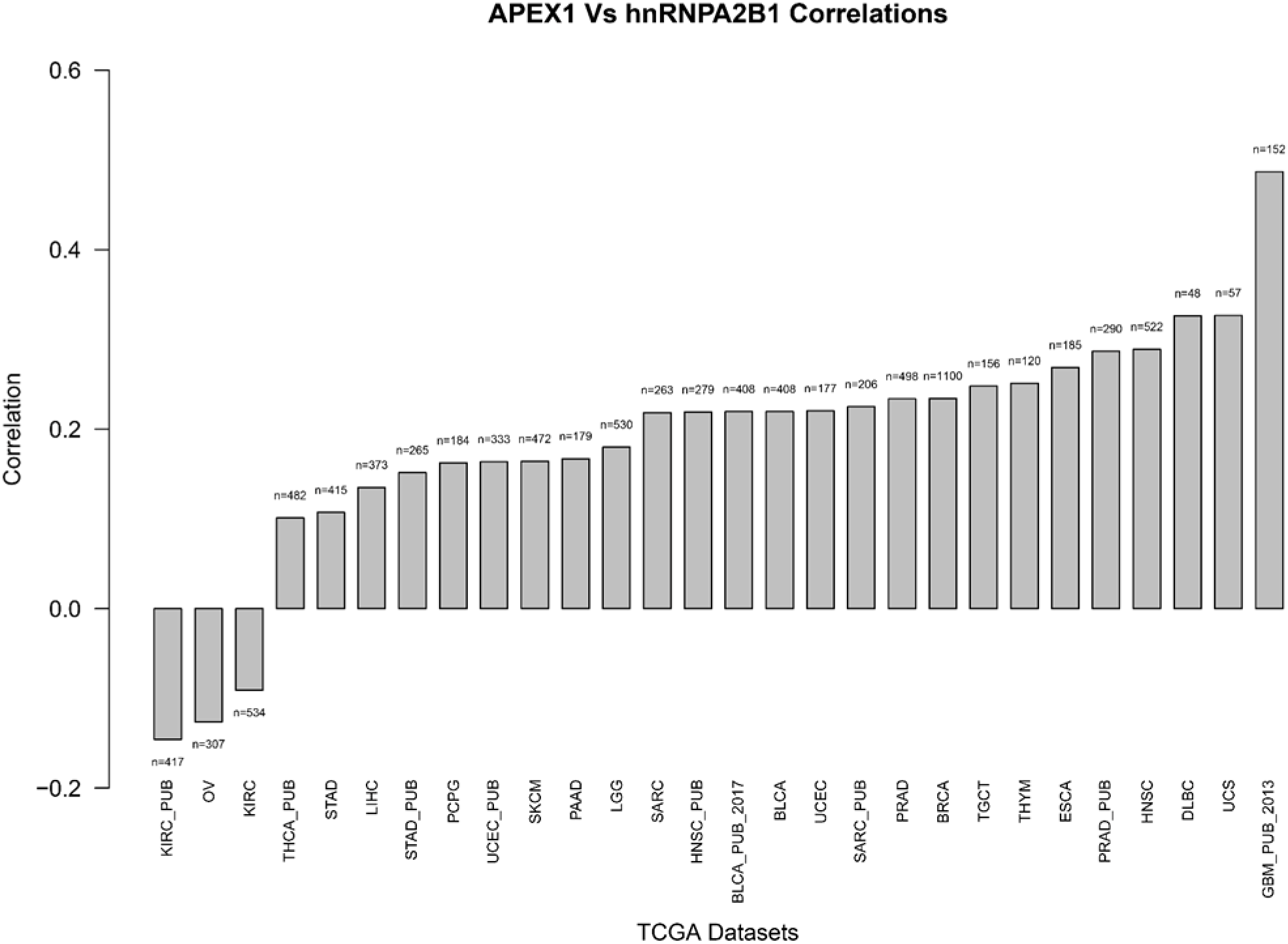
APE1 Vs hnRNPA2B1 correlation analysis was carried out consulting 44 TCGA datasets. The histograms show significant trend of correlation in the various cancer datasets analyzed.

### Supplementary Table Legends

**Supplementary Table 1**: Differentially Expressed Features (DEFs). In DEFs Summary (Sheet 1) for each comparison, the total number of DEFs is shown together with the percentage and total number of miRNAs and other non-coding RNAs in up-regulation or down-regulation. In the other sheets, for each comparison, complete list of DEFs.

**Supplementary Table 2**. Table of miRNA gene targets for each of the performed comparisons. In particular, for each comparison, we have considered in this table only the up-regulated miRNAs. The target identification was performed using the target prediction tool of QIAGEN Ingenuity Pathway Analysis (IPA) filtering for only experimentally validated miRNAs.

**Supplementary Table 3**. Table of miRNA gene targets for each of the performed comparisons. In particular, for each comparison, we have considered in this table only the down-regulated miRNAs. The target identification was performed using the target prediction tool of QIAGEN Ingenuity Pathway Analysis (IPA) filtering for only experimentally validated miRNAs.

**Supplementary Table 4:**
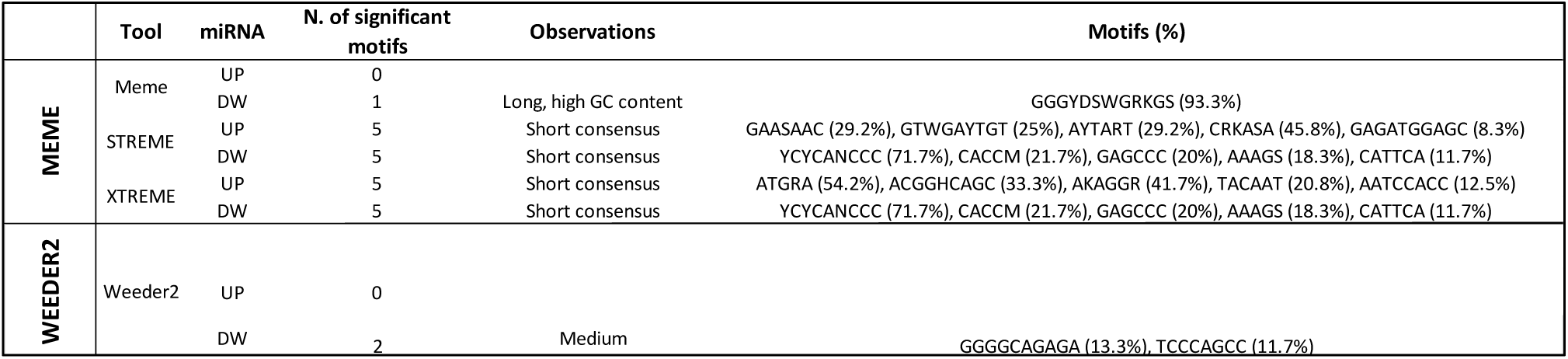
Table of performed analysis with MEME suite and Weeder2 for both up-regulated and down-regulated miRNAs. The number of statistically significant motif is shown together with potential observations and the motif and relative percentage found to be enriched in the specific miRNA subset.

**Supplementary Table 5. Identification of meme_motif1.** Table of 56 down-regulated miRNAs possessing GGGYDSWGRKGS motif and relative *p-value*, which represents the statistical significance of the observed motif of interest. The analysis was performed by making use of the tool MEME, belonging to the MEME suite.

**Supplementary Table 6**. Table of enriched motifs occurrences in the EVDW subset for each of the miRNAs. Ratio of average motif count in EVDW in respect to EVBG are reported. P-value was calculated for each motif with the motifcounter R package. For every miRNA of the subset, hairpin sequence and the corresponding available annotation is reported. The number of motif occurrences for each miRNA are coloured in gradient from white (0 occurrences) to dark green (5 occurrences).

**Supplementary Table 7:**
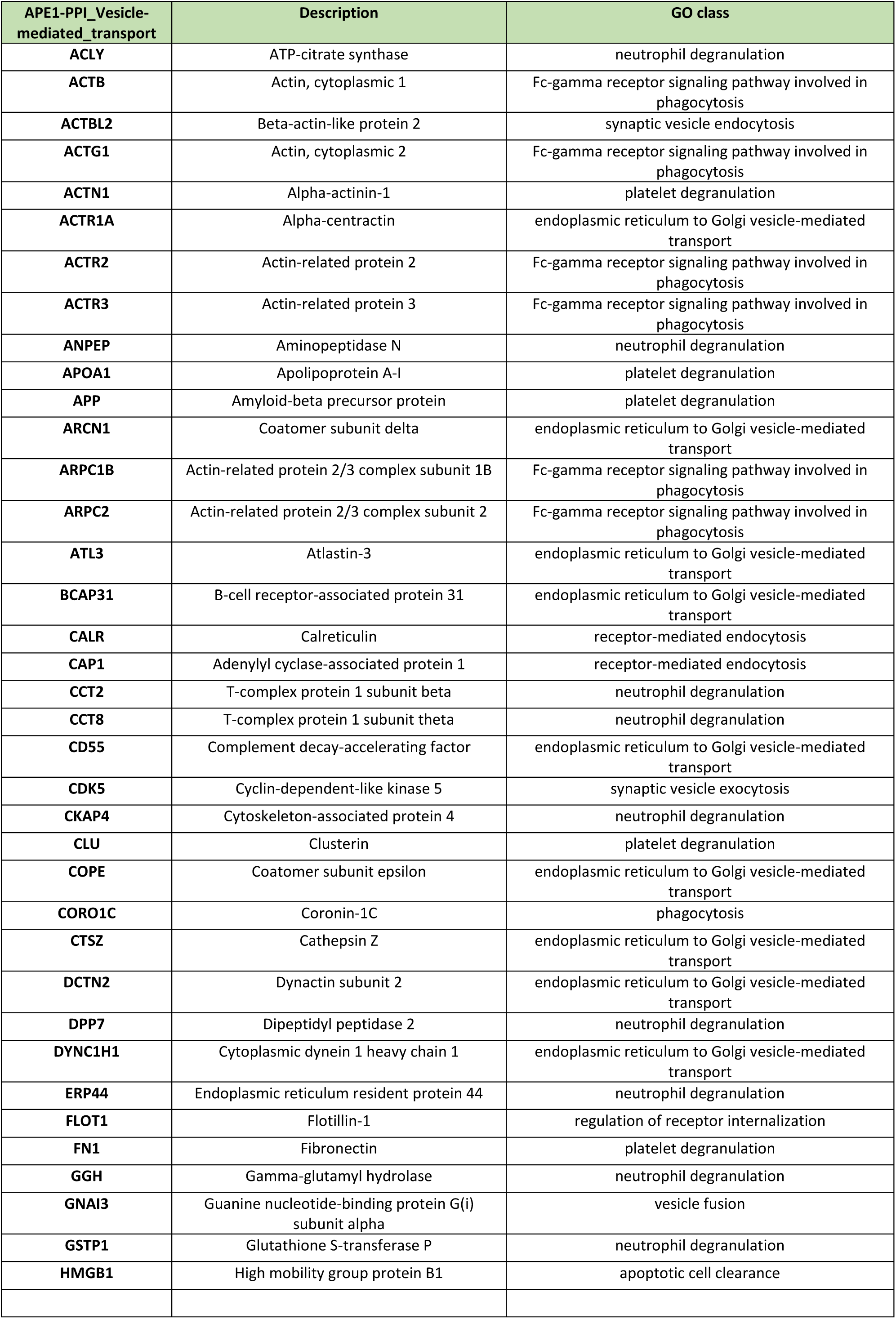
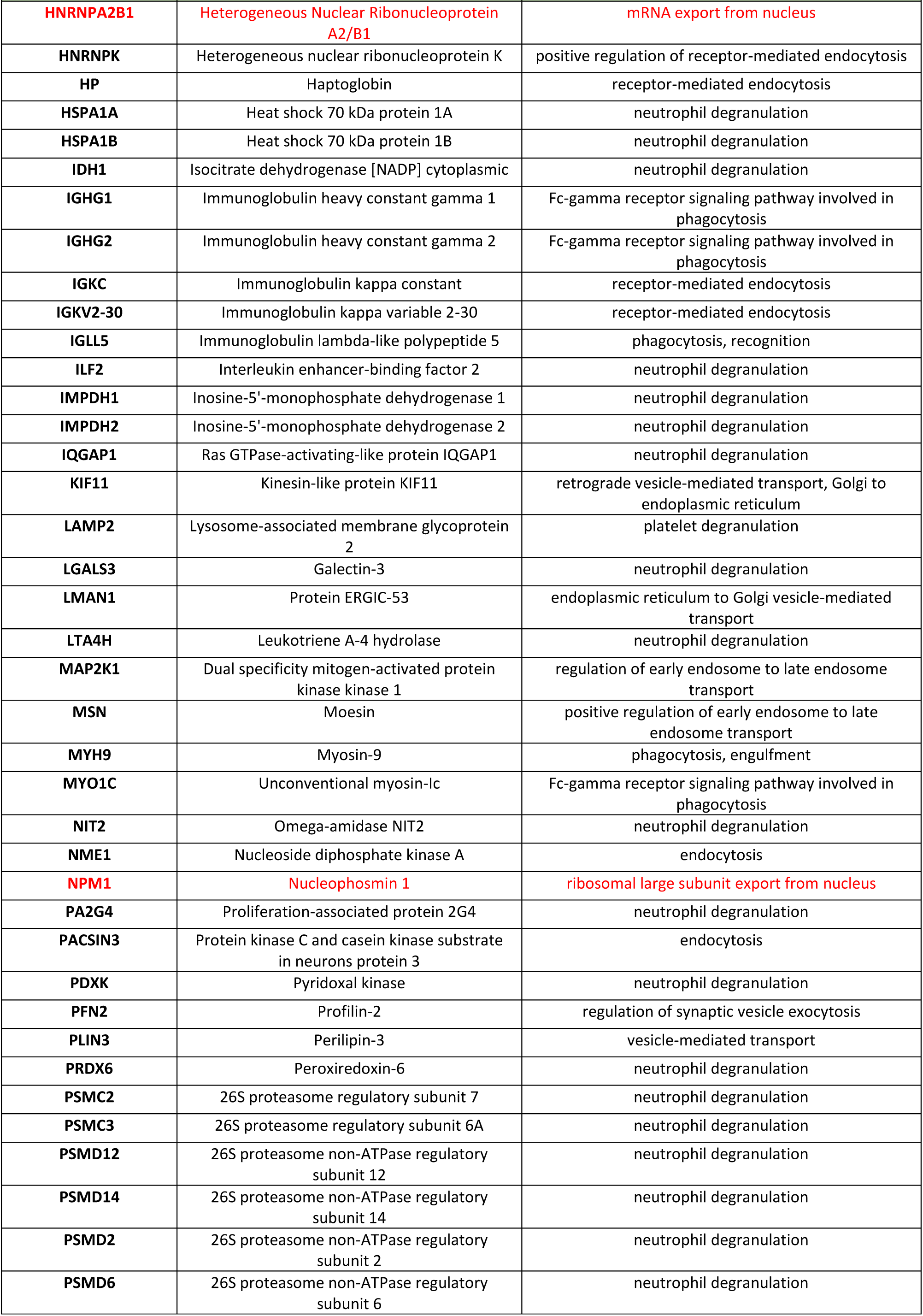
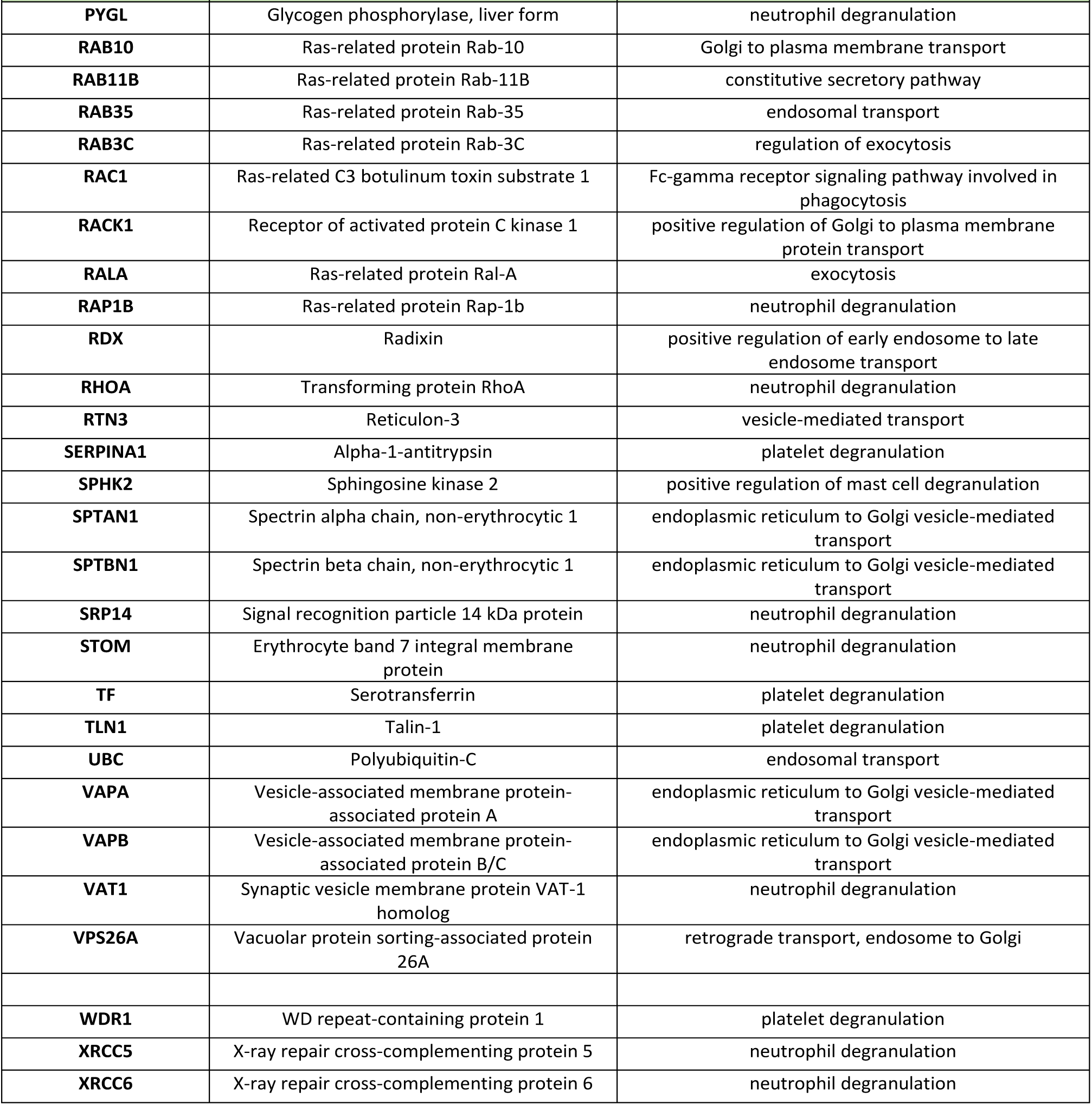
APE1 protein interactors involved in vesicles-mediated transport. In red protein selected for the validation of the interplay with APE1 for the regulation of miRNA sorting in this study.

**Supplementary Table 8**: Motif discovery analysis of the BGVS and BCCD EXO-motifs in four datasets of dysregulated miRNAs (DE-miRNAs) profiled in APE1 knocked down cells or oxidative stress conditions. The FIMO program was used to identify the presence of two EXO-motif associated with EVs secretion of mature miRNAs. motif1: BGVS ([GUC]G[ACG][GC]); motif2: BCCD ([GUC]CC[UGA]). Data show the occurrences of motif1 and motif2, respectively, in JHH-6 (worksheets a-b), A549 (worksheets c-d) and HeLa cells (APE1-depleted (worksheets e-f) and H2O2-treated (worksheets g-h)).

**Supplementary Table 9:** Number of identified occurrences of the two EXO-motifs associated with miRNAs sorting in EVs of the four examined cell lines.

|  | motif1 only | motif2 only | both |
| --- | --- | --- | --- |
| JHH-6 DE-miRNAs | 2 | 2 | 2 |
| A549 DE-miRNAs | 16 | 14 | 20 |
| H2O2 HeLa DE-miRNAs | 10 | 11 | 10 |
| siAPE1 HeLa DE-miRNAs | 16 | 13 | 29 |

**Supplementary Table 10**. **Analysis of miR-1246 validated targets.** List of the twelve validated targets of miR-1246 (TarBase v8.0) differentially expressed upon loss of APE1 expression in HeLa cells (fold change ≥ 1.5, q-value ≤ 0.05). Targets are sorted by decreasing fold change.

**Supplementary Table 11:**
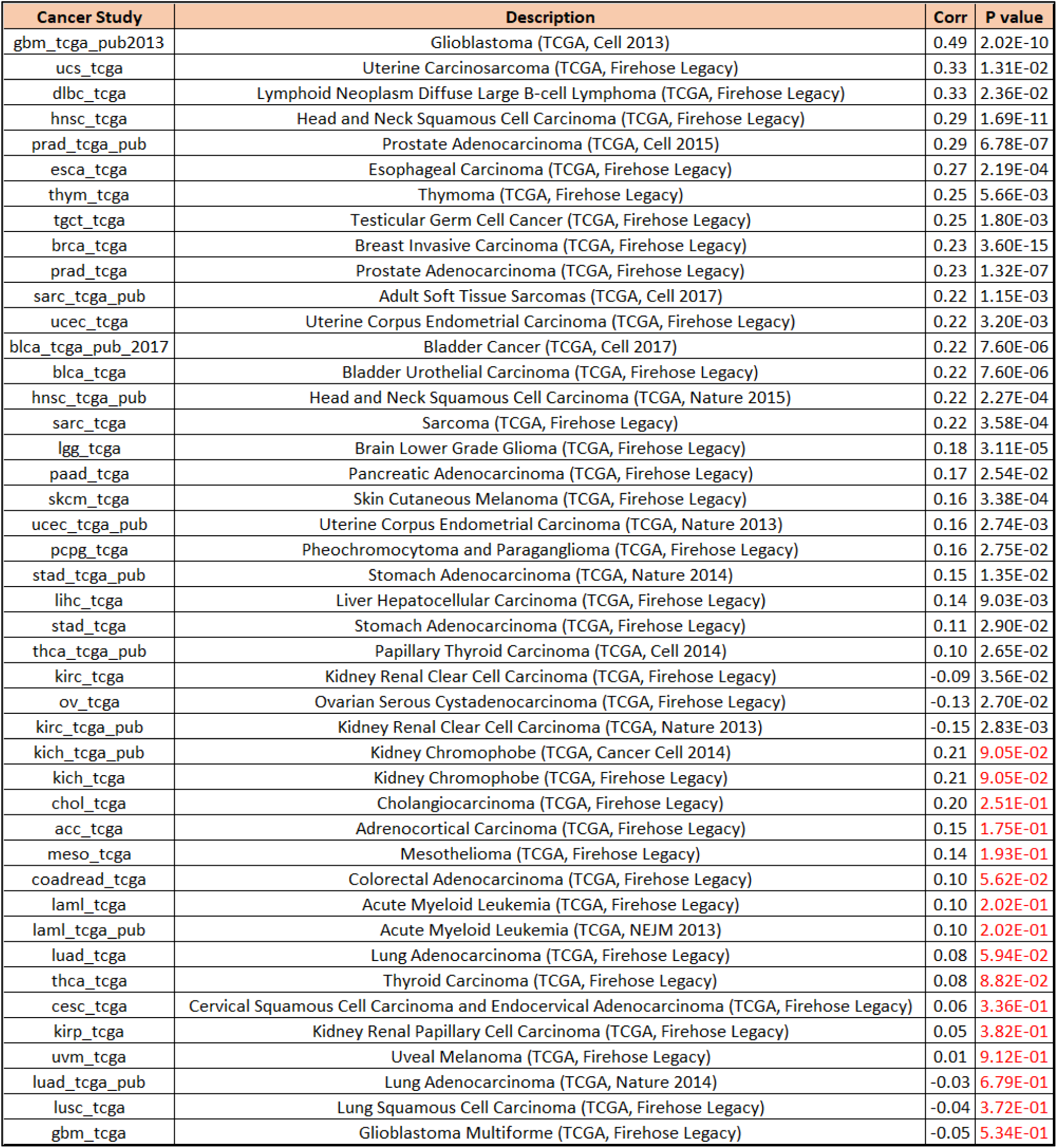
Correlation analysis of the gene expression levels of APE1 and hnRNPA2B1 in the examined TCGA data sets. The Spearman correlation values and the associated P value are shown, red indicating statistically not significant results.

| Cancer Study | Description | Corr | P value |
| --- | --- | --- | --- |
| gbm_tcga_pub2013 | Glioblastoma (TCGA, Cell 2013) | 0.49 | 2.02E-10 |
| ucs_tcga | Uterine Carcinosarcoma (TCGA, Firehose Legacy) | 0.33 | 1.31E-02 |
| dlbc_tcga | Lymphoid Neoplasm Diffuse Large B-cell Lymphoma (TCGA, Firehose Legacy) | 0.33 | 2.36E-02 |
| hnsc_tcga | Head and Neck Squamous Cell Carcinoma (TCGA, Firehose Legacy) | 0.29 | 1.69E-11 |
| prad_tcga_pub | Prostate Adenocarcinoma (TCGA, Cell 2015) | 0.29 | 6.78E-07 |
| esca_tcga | Esophageal Carcinoma (TCGA, Firehose Legacy) | 0.27 | 2.19E-04 |
| thym_tcga | Thymoma (TCGA, Firehose Legacy) | 0.25 | 5.66E-03 |
| tgct_tcga | Testicular Germ Cell Cancer (TCGA, Firehose Legacy) | 0.25 | 1.80E-03 |
| brca_tcga | Breast Invasive Carcinoma (TCGA, Firehose Legacy) | 0.23 | 3.60E-15 |
| prad_tcga | Prostate Adenocarcinoma (TCGA, Firehose Legacy) | 0.23 | 1.32E-07 |
| sarc_tcga_pub | Adult Soft Tissue Sarcomas (TCGA, Cell 2017) | 0.22 | 1.15E-03 |
| ucec_tcga | Uterine Corpus Endometrial Carcinoma (TCGA, Firehose Legacy) | 0.22 | 3.20E-03 |
| blca_tcga_pub_2017 | Bladder Cancer (TCGA, Cell 2017) | 0.22 | 7.60E-06 |
| blca_tcga | Bladder Urothelial Carcinoma (TCGA, Firehose Legacy) | 0.22 | 7.60E-06 |
| hnsc_tcga_pub | Head and Neck Squamous Cell Carcinoma (TCGA, Nature 2015) | 0.22 | 2.27E-04 |
| sarc_tcga | Sarcoma (TCGA, Firehose Legacy) | 0.22 | 3.58E-04 |
| lgg_tcga | Brain Lower Grade Glioma (TCGA, Firehose Legacy) | 0.18 | 3.11E-05 |
| paad_tcga | Pancreatic Adenocarcinoma (TCGA, Firehose Legacy) | 0.17 | 2.54E-02 |
| skcm_tcga | Skin Cutaneous Melanoma (TCGA, Firehose Legacy) | 0.16 | 3.38E-04 |
| ucec_tcga_pub | Uterine Corpus Endometrial Carcinoma (TCGA, Nature 2013) | 0.16 | 2.74E-03 |
| pcpg_tcga | Pheochromocytoma and Paraganglioma (TCGA, Firehose Legacy) | 0.16 | 2.75E-02 |
| stad_tcga_pub | Stomach Adenocarcinoma (TCGA, Nature 2014) | 0.15 | 1.35E-02 |
| lihc_tcga | Liver Hepatocellular Carcinoma (TCGA, Firehose Legacy) | 0.14 | 9.03E-03 |
| stad_tcga | Stomach Adenocarcinoma (TCGA, Firehose Legacy) | 0.11 | 2.90E-02 |
| thca_tcga_pub | Papillary Thyroid Carcinoma (TCGA, Cell 2014) | 0.10 | 2.65E-02 |
| kirc_tcga | Kidney Renal Clear Cell Carcinoma (TCGA, Firehose Legacy) | -0.09 | 3.56E-02 |
| ov_tcga | Ovarian Serous Cystadenocarcinoma (TCGA, Firehose Legacy) | -0.13 | 2.70E-02 |
| kirc_tcga_pub | Kidney Renal Clear Cell Carcinoma (TCGA, Nature 2013) | -0.15 | 2.83E-03 |
| kich_tcga_pub | Kidney Chromophobe (TCGA, Cancer Cell 2014) | 0.21 | 9.05E-02 |
| kich_tcga | Kidney Chromophobe (TCGA, Firehose Legacy) | 0.21 | 9.05E-02 |
| chol_tcga | Cholangiocarcinoma (TCGA, Firehose Legacy) | 0.20 | 2.51E-01 |
| acc_tcga | Adrenocortical Carcinoma (TCGA, Firehose Legacy) | 0.15 | 1.75E-01 |
| meso_tcga | Mesothelioma (TCGA, Firehose Legacy) | 0.14 | 1.93E-01 |
| coadread_tcga | Colorectal Adenocarcinoma (TCGA, Firehose Legacy) | 0.10 | 5.62E-02 |
| laml_tcga | Acute Myeloid Leukemia (TCGA, Firehose Legacy) | 0.10 | 2.02E-01 |
| laml_tcga_pub | Acute Myeloid Leukemia (TCGA, NEJM 2013) | 0.10 | 2.02E-01 |
| luad_tcga | Lung Adenocarcinoma (TCGA, Firehose Legacy) | 0.08 | 5.94E-02 |
| thca_tcga | Thyroid Carcinoma (TCGA, Firehose Legacy) | 0.08 | 8.82E-02 |
| cesc_tcga | Cervical Squamous Cell Carcinoma and Endocervical Adenocarcinoma (TCGA, Firehose Legacy) | 0.06 | 3.36E-01 |
| kirp_tcga | Kidney Renal Papillary Cell Carcinoma (TCGA, Firehose Legacy) | 0.05 | 3.82E-01 |
| uvm_tcga | Uveal Melanoma (TCGA, Firehose Legacy) | 0.01 | 9.12E-01 |
| luad_tcga_pub | Lung Adenocarcinoma (TCGA, Nature 2014) | -0.03 | 6.79E-01 |
| lusc_tcga | Lung Squamous Cell Carcinoma (TCGA, Firehose Legacy) | -0.04 | 3.72E-01 |
| gbm_tcga | Glioblastoma Multiforme (TCGA, Firehose Legacy) | -0.05 | 5.34E-01 |

### Supplementary Methods

#### Bioinformatics analyses

We used sRNAtoolbox, a collection of small RNA analysis tools, to generate the features’ counts. The generated feature counts were consequently analysed using R’s package DESeq2. The normalized count matrix (obtained from variance stabilizing transformation (VST) method as implemented in DESeq2 package) was used to explore high-dimensional data property with Principal Component Analysis (PCA) coupled with a dimensionality reduction algorithm used in the DESeq2 package. Differentially expressed features (DEFs) were selected with a p-adjusted cut off of 0.05 and a log2 Fold Change value greater than 1 (up-regulated DEFs) or lower than -1 (down-regulated DEFs). P-value was adjusted for multiple testing using the Benjamini–Hochberg (BH) correction with a false discovery rate (FDR) ≤ 0.05. DEFs were then analyzed with a hierarchical clustering method, using correlation distance. Visualization of log2-normalized values and clustering was obtained thanks to pheatmap package, while visualization of DEGs in volcano plots were acquired using the EnhancedVolcano package. Functional annotation was performed for all the comparisons and for feature list of interest. We used both QIAGEN Ingenuity Pathway Analysis using standard parameters and g:Profiler, whose results were visualized with ggplot2 and gplots R packages.

Among all the classes of molecular biotypes, accordingly to the ENSEMBL annotation database, we selected those classified as miRNA obtaining 68 down-regulated miRNAs in EV comparison (EVDW gene list). Hairpin sequences were retrieved from MiRBase website obtaining 62 unique precursors from EVDW gene list (accessed on October 2023).

Our next aim was focusing on analysing the occurrences of the coreExomotif, usually consisting of 4-nucleotides sequences. In order to be unbiased, we searched all possible four-nucleotide short sequences (n=256) both in the precursors and in a list of not differentially expressed miRNAs as background (EVBG, miRNAs n=318, unique precursors n=265).

To obtain EVBG miRNAs, they had to satisfy the following three criteria: a) the absolute value of log2 fold change had to be less than 1; b) the p-adjusted had to be greater than 0.05; c) the mean log2 normalized expression greater than 7. This filtering was performed in order to retrieve miRNAs that were not differentially expressed but were present in both condition of the comparison, to avoid selecting miRNAs called as not differentially expressed simply because they were not present. Using R/Bioconductor environment we counted each motif in every precursor and to calculated the enrichment and their p-values, we used motifEnrichment function (package motifcounter, (10)). To compare the difference in terms of motif counts between EVDW in respect to EVBG, we performed Wilcoxon rank test. In particular, we calculated and compared the average count of each motif normalized by the total number of precursor in each set.

EVspecificDW gene list was obtained selecting the down-regulated mature DE-miRNAs upon APE1 silencing in EV comparison, that were also not differentially expressed in cellular comparison (using the thresholds described above). We conducted for this set in respect to EVBG the Wilcoxon rank test.

coreEXO-motif (n=11) was derived from (11).

Heatmap was generated using complexHeatmap package (12).

#### Survival analysis

Survival were performed as reported in (4).

#### Gene Ontology analysis

We queried the AmiGO 2 (http://amigo.geneontology.org/amigo/landing) and the BioCyc (https://biocyc.org/) websites (last accessed: April 2020) to find all the human proteins associated with the “vesicle-mediated transport” term (GO:0016192). We obtained 4587 and 301 hits in AmiGO 2 and BioCyc, respectively, corresponding to 2181 and 298 unique human proteins. We merged this data to obtain the final dataset (n=2208) of non-redundant human proteins that was used for the downstream analysis.

#### EXO-motifs discovery in JHH-6, A549 and HeLa mature DE-miRNAs

The two EXO-motifs consensus sequences were recovered from (3) and summarized as motif1: BGVS ([GUC]G[ACG][GC]) and motif2: BCCD ([GUC]CC[UGA])). We downloaded from miRBase (ftp://mirbase.org/pub/mirbase/CURRENT/mature.fa.gz) the FASTA formatted mature miRNA sequences. We generated four subsets corresponding to the human mature miRNAs that were differentially expressed in APE-1 depleted JHH-6, A549 and HeLa cells and in H2O2-treated HeLa cells. We repeated the same procedure for the nineteen miRNAs regulated by mutNPM1, generating a fifth dataset. We used the MEME-Suite program FIMO (http://meme-suite.org/tools/fimo) to scan these five sets of FASTA sequences to find individual matches to each of the two EXO-motifs, with default settings except for: Match p-value < 0.1. One output file was generated for each EXO-motif/cell-type combination.

#### miRNA targets functional enrichment analysis

DE-miRNAs human validated targets were retrieved from TarBase v.8 (13). Functional enrichment analysis was performed on the identified unique target genes (n=1501) with the Cytoscape (14) plugin ClueGO (15) to identify functionally enriched terms. The following functional databases were queried: CLINVAR_Human-diseases (08.05.2020), CORUM_CORUM-3.0-FunCat-MIPS (03.09.2018), GO_BiologicalProcess (03.05.2021), GO_ImmuneSystemProcess (03.05.2021), KEGG (03.05.2021), KEGG-HUMAN-DISEASE (03.05.2021), REACTOME_Pathways (03.05.2021) and WikiPathways (08.05.2020). Default parameters were applied except for “Number of Genes” = 70 and “Min Percentage” = 12.5. A two-sided hypergeometric test was used to determine the probability that each functional term was assigned to the gene sets due to chance alone.

#### Western blot analysis

Protein extracts were quantified with the Bradford solution (Biorad, California, USA), according to the manufacturer’s instructions, loaded in 10 or 12% gel (SureCast Acrylamide Solution (40%), Invitrogen), and transferred into nitrocellulose membrane (Amersham, Little Chalfont, UK). Western blot analyses were executed using the listed antibodies: anti-APE1 (13B8E5C2, Novus), anti-β-tubulin (T0198, Sigma-Aldrich), anti-FLAG M2 (F1804, Sigma Aldrich), anti-hnRNPA2B1 (PA5-34939, Thermo Fischer Scientific), anti-NPM1 (ab15440, Abcam), anti-Syntenin (EPR8102) (ab133267, Abcam). After incubation with primary antibodies, membranes were washed three times with PBS 0.1% Tween-20, (Sigma Aldrich), incubated for 1 h at room temperature with the appropriate IRDye800/IRDye600 labelled secondary antibodies (diluted to 1:2000). The acquisition of the images and the quantifications analyses were achieved using the Odyssey CLx Infrared Imaging system (LI-COR GmbH, Germany).

#### Co-immunoprecipitation

On day one 3*10^6 JHH-6 cells were seeded in 10 cm dish and the day after were transfected with the FLAG-tagged APE1 protein encoding plasmid using the Lipofectamine 2000 reagent (Invitrogen, California United States), according to the manufacturer’s instructions. The following day, cells were washed twice with PBS, and resuspended in eight hundred μl of lysis buffer (50 mM Tris HCl pH 7.4, 150 mM NaCl, 1 mM EDTA, 1% Triton X-100, supplemented with 1 mM protease inhibitor cocktail (Sigma), 0.5 mM phenylmethylsulfonyl fluoride (PMSF), 1 mM NaF, 1 mM Na3VO4). The lysis was performed under rotation for 20 minutes at 4°C. The lysates were then harvested and centrifuged at 12,000 x *g* for 10 min at 4°C. Supernatants were collected and 50μl of lysate for each condition were kept as INPUT and quantified with the Bradford solution. In parallel, co-immunoprecipitation was carried out with the anti-FLAG M2 affinity gel (Sigma) for 3 hr on the rocker at 4°C. After incubation, the resin was centrifuged at 8,000 x *g* for 1 min, washed three times with Tris-buffered saline (TBS) and then the elution was performed with 0.15 mg ml-1 of FLAG peptide (Sigma Aldrich, Milan, Italy) in TBS complemented with protease inhibitors for 30 min at 4°C. Coomassie brilliant blue staining loading 3μl of IP in a 10% polyacrylamide gel was executed to normalize the different IPs.

The treatment with RNaseA (Sigma) was performed at the concentration of 100μg/ml for 30 minutes at 37°C. For treatments with DNaseI RNase free (Thermo Fisher Scientific), lysates were prepared using the following lysis buffer: DNaseI Reaction buffer, protease inhibitor cocktail (Sigma), 25 mM NaCl, 1% Triton X-100. Once lysates were ready, DNase I was added to the lysate and the treatment was performed for 30 minutes at 37°C. The inactivation of DNaseI was executed simply adding DTT 1mM to the lysate. No heat inactivation was made to avoid protein degradation.

#### siRNA silencing and miRNA extraction

3.0*10^^6^ JHH-6 cells were seeded in 150 cm dishes. The day after, cells were transfected with 17 pmol of custom hAPE1 siRNA (Sense: 5’ U.A.C.U.C.C.A.G.U.C.G.U.A.C.C.A.G.A.C.C.U.U.U 3’; Antisense: 5’ A.G.G.U.C.U.G.G.U.A.C.G.A.C.U.G.G.A.G.U.A.U.U 3’) or with non-targeting siRNA pool (siSCR) as a control, (GE Healthcare Dharmacon, Colorado, Unites States). The transfection was carried out with the DharmaFECT transfection reagent (Thermo Fisher Scientific, Massachusetts, United States), according to the manufacturer’s instructions. Six hr after transfections, fresh medium was added to the complexes for reducing cytotoxicity. The day after transfection, cells were washed twice with PBS, medium supplemented with 10% FBS exosome-depleted (Thermo Fisher Scientific), 2 mM L-glutamine, 100 U/ml penicillin, 100 μg/ml streptomycin (Euroclone) was added and after 72 hr from transfection cells and media were collected. Cellular and exosomal miRNAs extraction was executed, using respectively the miRNeasy Mini Kit (QIAGEN, Hilden, Germany), and the Total Exosome RNA & Protein Isolation Kit (Thermo Fisher Scientific) according to the manufacturer’s instructions.

#### RNA immunoprecipitation (RIP)

RNA immunoprecipitation were performed as reported in (6).

#### Total miRNAs quantification and quantitative reverse Transcriptase-PCR (qRT-PCR)

Cell and EXO miRNAs were quantified using the Qubit™ microRNA Assay Kit according to the manufacturer’s instructions and the Qubit 4 Fluorometer instrument (Thermo Fisher Scientific). miRNA expression profiles were evaluated by RT-qPCR using the TaqMan Advanced miRNA assay (Life Technologies, Carlsbad, CA, USA). Briefly, 1.5 ng of miRNAs were retro-transcribed using the TaqMan® Advanced miRNA cDNA Synthesis Kit (Life Technologies) according to the manufacturer’s instructions. qRT-PCR reactions were carried out using the TaqMan Fast Advanced Master Mix, according to the manufacturer’s instructions and the qRT-PCR were performed using the CFX Touch™ Real-Time PCR System (Bio-Rad, Hercules, CA). For endogenous microRNAs RT-qPCR, the analyses were obtained applying the ΔΔct method, using the expression of miR-16-5p as reference. For Exo miRNAs analyses, a quantitative normalisation was carried out retro-transcribing the same amount of microRNAs for each analysed condition.

For the analysis of the pri-mRNAs expression, 1 µg of total RNA was reverse transcribed using the SensiFAST cDNA synthesis kit (Meridian Bioscience, Ohio, USA), according to the manufacturer’s instructions and qRT-PCR reactions and analyses were carried out as indicated above.

#### List of Taqman probes used

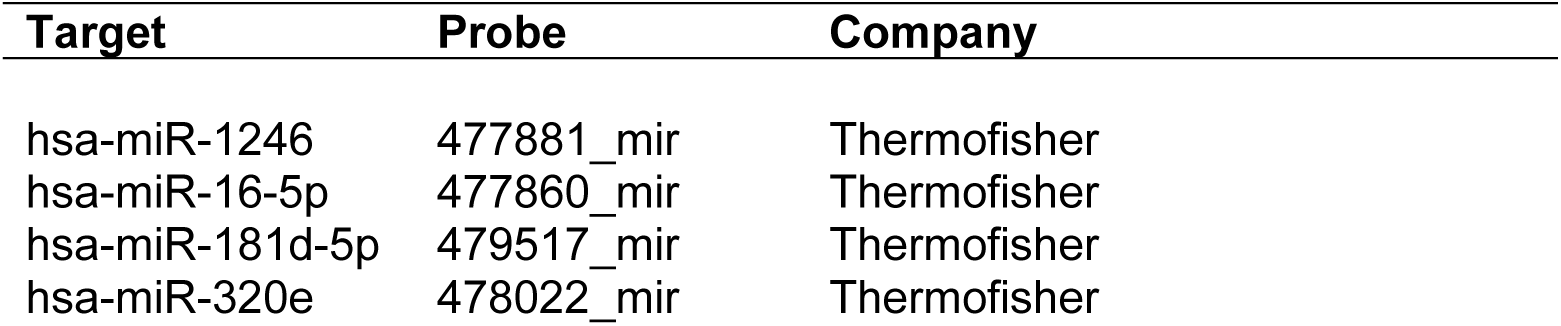

#### Recombinant proteins production

The constructs pGex3X, pGex3X-hAPE1, pGex3X-N33hAPE1, pGEX-6P-1-HNRNPA2B1, coding for the glutathione S-transferase (GST) protein tag, rGST APE1^WT^, rGST APE1^NΔ33^ and rGST hnRNPA2B1 were used to transform *E. coli* BL21(DE3). Protein expression was induced with 1 mM (or 0.2 mM only for hnRNPA2B1) isopropyl-b-D-thiogalactopyranoside (IPTG), and then purified on an AKTA Prime FPLC system (GE Healthcare) by using a GSTrap HP column (GE Healthcare). Quality of purification control and protein quantification were performed by Coomassie Brilliant Blue (Sigma, Milan, Italy) staining of SDS–PAGE gels.

#### List of RNA probes used for EMSA experiments

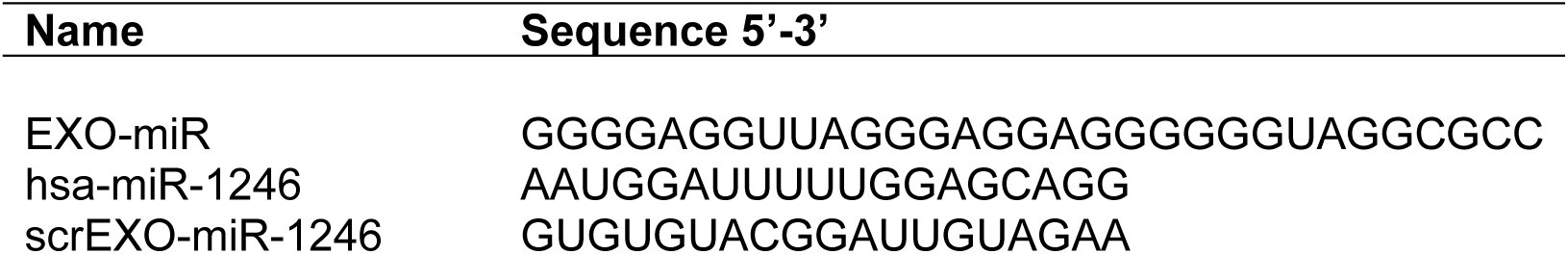

RNA probes listed in the chart were purchased from Metabion, Germany.

#### Electrophoretic Mobility Shift Assay (EMSA)

EMSA assays were carried out as explained in previous work from our research group (16). Recombinant proteins were incubated with RNA probe (pretreated with DTT 1 mM at 70 °C for 2 min) for 20 min at room temperature (RT) before loading on gel. When two proteins were loaded together with the probe, they were incubated for 20 min at RT prior to the RNA probe addition, in order to allow the protein-protein interaction to occur. The same pre-incubation step was added also when using the inhibitors of RNA-protein interaction. Samples were prepared with the use of EMSA buffer (20 mM Hepes pH 7.5, 50 mM KCl, 0.5 μg/ml BSA, 0.25% glycerol). EMSA run was performed with pre-chilled 0.5X Tris/Borate/EDTA buffer (TBE) at 4 °C. Images were acquired by using an Odyssey CLx Infrared Imaging system (Li-Cor Biosciences).

## Notes

https://www.ncbi.nlm.nih.gov/geo/query/acc.cgi?acc=GSE230874

### Competing Interest Statement

The authors have declared no competing interest.

